# Whole-brain analyses identify anterior cingulate μ-opioid signaling as a critical mediator of placebo analgesia in neuropathic pain

**DOI:** 10.64898/2026.06.04.729974

**Authors:** Simran K. Rehal, Damien C. Boorman, Chulmin Cho, Seyed Asaad Karimi, Maryam I. Fazili, Cesar AO. Coelho, Jacob Burek, Farhada Khaled, Shivani R. Gami, Paul W. Frankland, Loren J. Martin

**Affiliations:** Department of Cell and Systems Biology, University of Toronto; Toronto, Ontario, M5S 3G5, Canada; Department of Psychological and Brain Sciences, University of Toronto Mississauga; Mississauga, Ontario, L5L 1C6, Canada; Department of Psychology, University of Toronto, Toronto, ON M5S 3G3, Canada; Program in Neuroscience & Mental Health, Hospital for Sick Children, 555 University Avenue, Toronto, ON M5G 1X8, Canada; Department of Physiology, University of Toronto, Toronto, ON M5S 1A8, Canada; Child & Brain Development Program, Canadian Institute for Advanced Research (CIFAR), Toronto, ON M5G 1M1, Canada

## Abstract

Placebo analgesia reflects the capacity of learning and expectation to engage endogenous pain-control systems, yet the neural circuits that support this phenomenon remain poorly understood. Here, we establish a conditioning-based placebo analgesia paradigm in mice following peripheral nerve injury and combine behavioral assessment with whole-brain activity mapping and targeted circuit manipulations to define its neural substrates. Conditioning with morphine produced robust placebo analgesia, expressed as reduced sensory sensitivity in the absence of drug. Brain-wide mapping of c-Fos expression revealed distributed changes across cortical and subcortical regions accompanied by a reorganization of functional connectivity, consistent with coordinated network-level engagement. Network analyses identified shifts in hub structure and selective strengthening and weakening of inter-regional interactions during placebo analgesia. Causal manipulations demonstrated a critical role for the anterior cingulate cortex, with excitatory activation of this region blocking placebo analgesia, whereas inhibitory manipulations had no effect. In contrast, perturbation of other candidate regions, including the basomedial amygdala and paraventricular thalamus, did not alter placebo responses. Finally, selective targeting of μ-opioid receptor–expressing neurons in the anterior cingulate cortex revealed that this cell population is necessary for the expression of placebo analgesia. Together, these findings reveal a brain-wide reorganization of network interactions underlying placebo analgesia and identify a specific cortical opioid circuit that gates its expression.

**One-Sentence Summary:** Using a mouse model of nerve injury, this study shows that placebo analgesia arises from coordinated brain-wide network reorganization and is gated by μ-opioid receptor–expressing neurons in the anterior cingulate cortex.

## Introduction

Pain perception is fundamentally shaped by cognitive and contextual factors, enabling the same noxious stimulus to evoke dramatically different sensations depending on prior experience, expectation, and environmental cues[1, 2]. Placebo analgesia is one of the most compelling demonstrations of this cognitive-nociceptive interaction, in which cues previously associated with pain relief produce genuine analgesic effects in the absence of an active pharmacological intervention[3, 4]. The magnitude of these effects is clinically meaningful, as placebo responses account for an average of 20–40% of analgesic efficacy across studies[5] and, in controlled experimental settings, can produce analgesic effects comparable to those of low-dose morphine[6]. Placebo analgesia reflects the engagement of endogenous pain-modulatory systems[5, 7], yet the circuit architecture through which expectation and context recruit these systems remains incompletely defined.

Human neuroimaging and animal circuit studies have converged on a cortico-brainstem circuit for placebo analgesia. In humans, placebo analgesia recruits prefrontal, anterior cingulate, and insular cortices, together with brainstem regions including the periaqueductal gray (PAG), and reflects engagement of descending pain modulatory systems linking frontal–cingulate networks to PAG-centered control pathways[8, 9]. PET imaging has revealed changes in μ-opioid receptor binding within the anterior cingulate cortex (ACC), prefrontal cortex, and nucleus accumbens during placebo responses[10]. Complementing these findings, fMRI studies have shown a naloxone-sensitive coupling between the rostral ACC and the PAG that drives the recruitment of downstream brainstem structures, including the rostral ventromedial medulla (RVM), to mediate the behavioral expression of placebo analgesia[11]. In rodents, optogenetic and chemogenetic approaches have identified discrete circuits underlying conditioned placebo-like analgesia, including anterior cingulate projections to the pontine nucleus, which is both necessary and sufficient for analgesia[12], descending cortical–PAG pathways[13–15] and amygdala-dependent mechanisms that can support context-dependent analgesic learning[16]. However, these mechanistic insights derive nearly exclusively from acute pain models, leaving a gap in understanding how placebo analgesia operates under chronic pain conditions.

The transition from acute to chronic neuropathic pain involves central sensitization[17], enhanced descending facilitation from the rostral ventromedial medulla[18], diminished endogenous opioid function[19], and extensive structural and functional reorganization of cortical pain networks[20] — changes that may fundamentally alter the circuits through which placebo analgesia is expressed. While placebo hypoalgesia has been observed in some chronic pain populations, including patients with temporomandibular disorder[21], it remains unclear whether these effects engage the same cortico-brainstem analgesic circuits identified in acute pain models or arise through compensatory network reorganization. Given the widespread alterations in cortical and subcortical connectivity associated with chronic pain, placebo expression may depend on coordinated activity across distributed brain networks and therefore may be best understood at the network level.

Here, we used a morphine-conditioned placebo paradigm in mice with spared nerve injury (SNI) to test whether placebo analgesia can be expressed in a chronic neuropathic pain state and to define its underlying circuitry. Repeated morphine–context pairings were sufficient to condition placebo analgesia in SNI mice, and this response was abolished by μ-opioid receptor antagonism, confirming opioid-dependent placebo expression. Combining whole-brain c-Fos mapping with network analysis and causal manipulations, we find that placebo expression is not merely a suppression of pain-related activity but instead reflects a large-scale reorganization of brain-wide interactions, shifting from a predominantly subcortical organization toward greater cortical coordination. Within this reorganized network, the ACC emerged as a critical node, where reduced activity is permissive for placebo expression and local μ-opioid receptor signaling is required. Together, these results provide a circuit-level account of placebo analgesia in chronic pain that centers on cortical control mechanisms rather than recruitment of a single descending pathway.

## Results

### Nerve-injured mice exhibit morphine-conditioned placebo analgesia that is blocked by μ-opioid receptor antagonism

Pharmacological conditioning produces placebo responses in humans[22, 23] and rodents[24, 25], including morphine-conditioned analgesia in naïve mice[14]. Evidence in chronic pain models remains limited and inconsistent, with partial morphine-conditioned responses in neuropathic rats[26] and mixed outcomes following gabapentin conditioning[13, 27, 28]. Thus, we sought to determine whether robust, context-dependent placebo analgesia can be established in mice after nerve injury. Following spared nerve injury (SNI), mice underwent repeated morphine–context pairings, and analgesia was assessed on a saline test day in the same context (**Fig. 1a–c**). In all groups, morphine produced a consistent analgesic effect across conditioning days (Supplementary Fig. 1a,b). On the test day, conditioned mice (*Placebo*; Mor/Sal), which received morphine during conditioning but saline at test, showed a clear recovery in withdrawal thresholds relative to controls (*Natural history*; Sal/Sal), consistent with placebo analgesia. This effect was comparable to acute morphine analgesia (*Drug control*; Mor/Mor) and was abolished by naloxone (*Naloxone challenge*; Mor/Nlx), confirming opioid-dependent placebo expression (**Fig. 1d**). We next tested whether this response was mediated by specific opioid receptor subtypes. Systemic μ-opioid receptor antagonism with CTOP abolished the placebo response, whereas δ- and κ-opioid receptor antagonists had no effect (**Fig. 1e**).

**Figure 1.**
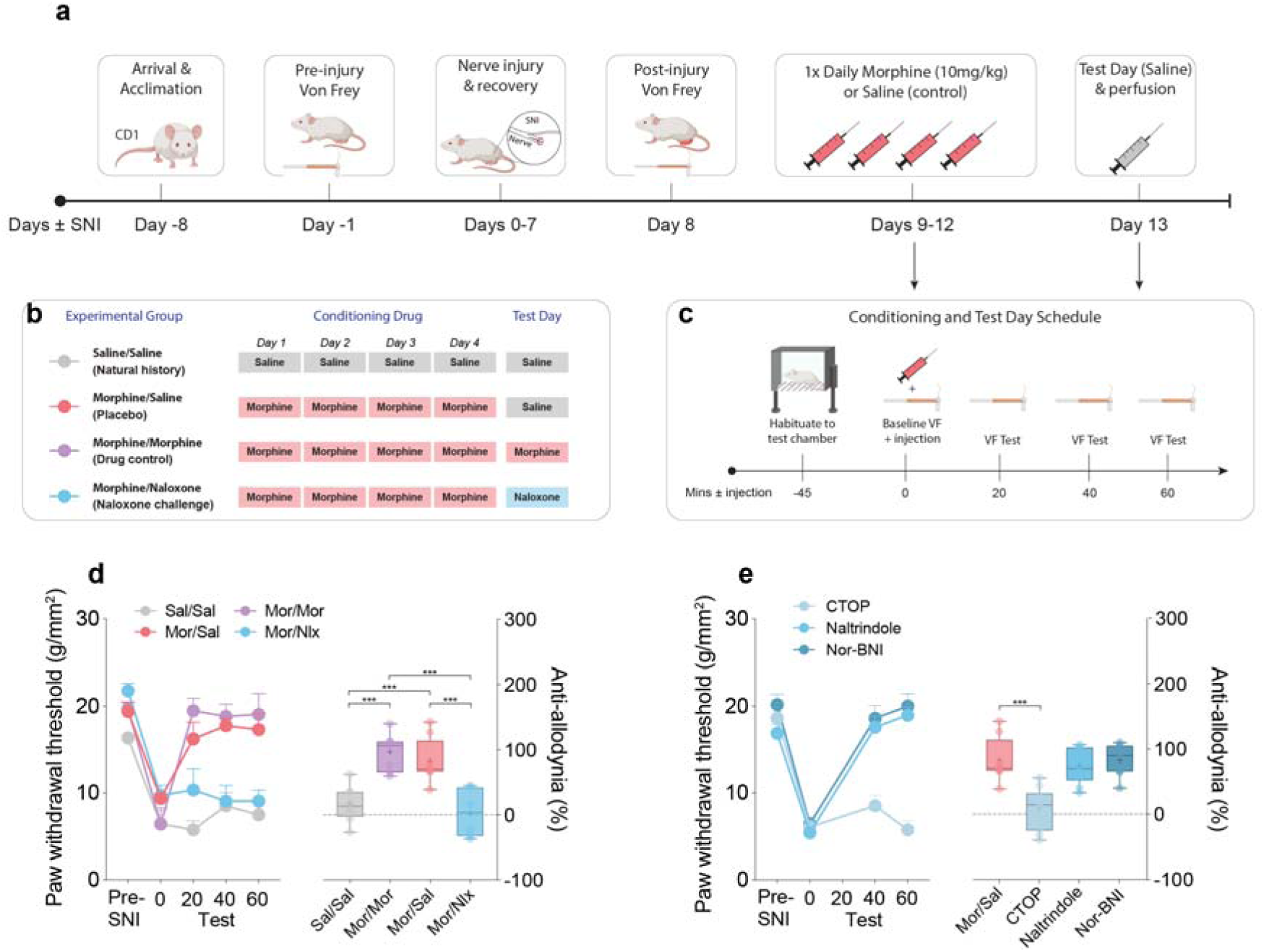
Experimental design and behavioral pharmacology of placebo analgesia in mice with SNI. **a** Schematic timeline of pharmacological conditioning procedure to elicit placebo analgesia in mice with spared nerve injury (SNI). CD1 mice were first acclimated and then underwent baseline von Frey testing prior to SNI surgery. 8 days post-surgery, mice underwent von Frey testing to establish a post-SNI baseline, followed by 4 days of conditioning with morphine (10 mg/kg, i.p.). On day 13 (Test Day), mice were given saline (placebo condition), von Frey tested, and then euthanized 90 minutes later and brains extracted. **b** Experimental groups and drug treatments across the conditioning and test phases, including repeated saline (Sal/Sal), morphine conditioning followed by saline on the test day (Mor/Sal), repeated morphine (Mor/Mor), and morphine conditioning followed by naloxone administration on the test day (Mor/Nlx). **c** Within-session testing procedure used during conditioning and placebo testing. **d** Withdrawal thresholds differed across treatment groups over time on the test day (group × time interaction, F(12,120)=6.23, p<0.0001), and anti-allodynia differed between groups (Welch’s ANOVA, F(3,14.22)=15.16, p=0.0001). Mor/Sal mice showed greater anti-allodynia than Sal/Sal controls (p=0.001), whereas naloxone blocked placebo expression (Mor/Sal vs Mor/Nlx, p=0.0032). Group sizes were Sal/Sal (n = 10), Mor/Sal (n = 8), Mor/Mor (n = 8), and Mor/Nlx (n = 8). **e** Opioid receptor antagonist testing showed a group × time interaction for withdrawal thresholds (F(6,81)=15.34, p<0.0001) and a group effect for anti-allodynia (Welch’s ANOVA, F(3,17.76)=11.97, p=0.0002). CTOP reduced placebo analgesia relative to saline-treated placebo mice (p=0.0019), whereas naltrindole and nor-BNI did not differ from saline treatment. Group sizes were CTOP (n = 10), naltrindole (n = 10), and nor-BNI (n = 10). Data shown as mean ± s.e.m. (line graphs) and individual data points with box-and-whisker plots. ***p<0.001, one-way ANOVA with post-hoc multiple comparisons with Tukey’s correction. Full statistical reporting is provided in Supplementary Data 1. Source data are provided as a Source Data file.

Having established morphine-conditioned placebo analgesia in neuropathic pain, we next tested whether altering conditioning parameters affected its expression. Reducing the conditioning dose of morphine (5 mg/kg) failed to yield significant placebo-associated anti-allodynia compared with the standard conditioning paradigm, whereas shortening the conditioning period to 2 days produced weaker and more variable placebo-associated anti-allodynia (Supplementary Fig. 2a–c). In contrast, conditioning with gabapentin (40 or 100 mg/kg) failed to elicit a placebo response despite producing robust analgesia during conditioning (Supplementary Fig. 2d,e). Placebo-associated anti-allodynia was observed in both male and female mice, with male mice showing slightly greater anti-allodynia than females (Supplementary Fig. 3a). Placebo responses were also observed across experiments performed by multiple blinded male and female experimenters with placebo magnitude varying across cohorts (Supplementary Fig. 3b). Placebo expression was nevertheless context-dependent, as dissociating the drug-paired and testing environments markedly reduced placebo-associated anti-allodynia (Supplementary Fig. 3c).

Given prior reports of conditioned analgesia in naïve rodents using thermal nociceptive assays, we also tested whether pharmacological conditioning could elicit placebo-associated analgesia in non-injured mice using the hot-plate test. Across experiments, we incorporated and progressively strengthened contextual and multisensory conditioning cues, including distinct conditioning chambers, visual and food-associated cues, and extended conditioning procedures. Despite robust morphine-induced thermal analgesia during conditioning, placebo-associated analgesia was not reliably observed on saline test days in naïve mice (Supplementary Fig. 4). Thus, conditioned placebo responses were not robustly expressed under our thermal testing parameters, potentially reflecting differences in contextual conditioning procedures and cue–assay pairing across studies[14, 29–33]

Finally, we tested whether interleaving morphine with saline alters tolerance development by comparing continuous morphine administration with a partial reinforcement schedule. Daily morphine administration produced a progressive loss of antinociceptive efficacy beginning around day 7, consistent with the development of tolerance[34]. In contrast, interleaving saline on days 5, 7, and 9 preserved within-session morphine responsiveness and attenuated tolerance development relative to continuous morphine exposure (Supplementary Fig. 5a,b). Consistent with this, intermittent morphine administration maintained higher antinociceptive efficacy on day 10, as assessed by % maximum possible effect (MPE), whereas continuous morphine administration produced marked reductions in morphine responsiveness (Supplementary Fig. 5c,d). This pattern aligns with dose-extending paradigms, in which interleaving active drug with placebo maintains drug-like responses through pharmacological conditioning[35, 36].

### Placebo analgesia produces region-specific changes across a distributed brain-wide network

To define brain-wide activity associated with placebo analgesia, we quantified the immediate early gene c-Fos across 56 regions spanning the cortex, limbic forebrain, thalamus, hypothalamus, and brainstem (**Fig. 2a**, Supplementary Fig. 6, and Supplementary Table 1). A home-cage control group was included to establish baseline activity across regions. Both natural history and placebo groups showed brain-wide increases in c-Fos expression compared to the home-cage control group. Compared to natural history, placebo expression was associated with widespread reductions in cortical activity alongside increased engagement of midbrain and brainstem structures involved in descending pain modulation (**Fig. 2b–d**). Bilateral reductions were observed across orbital frontal regions, lower-limb somatosensory and dorsal insular cortices, and both dorsal and ventral ACC subdivisions, consistent with reduced cortical engagement during placebo expression (**Fig. 2b,c**).

**Figure 2.**
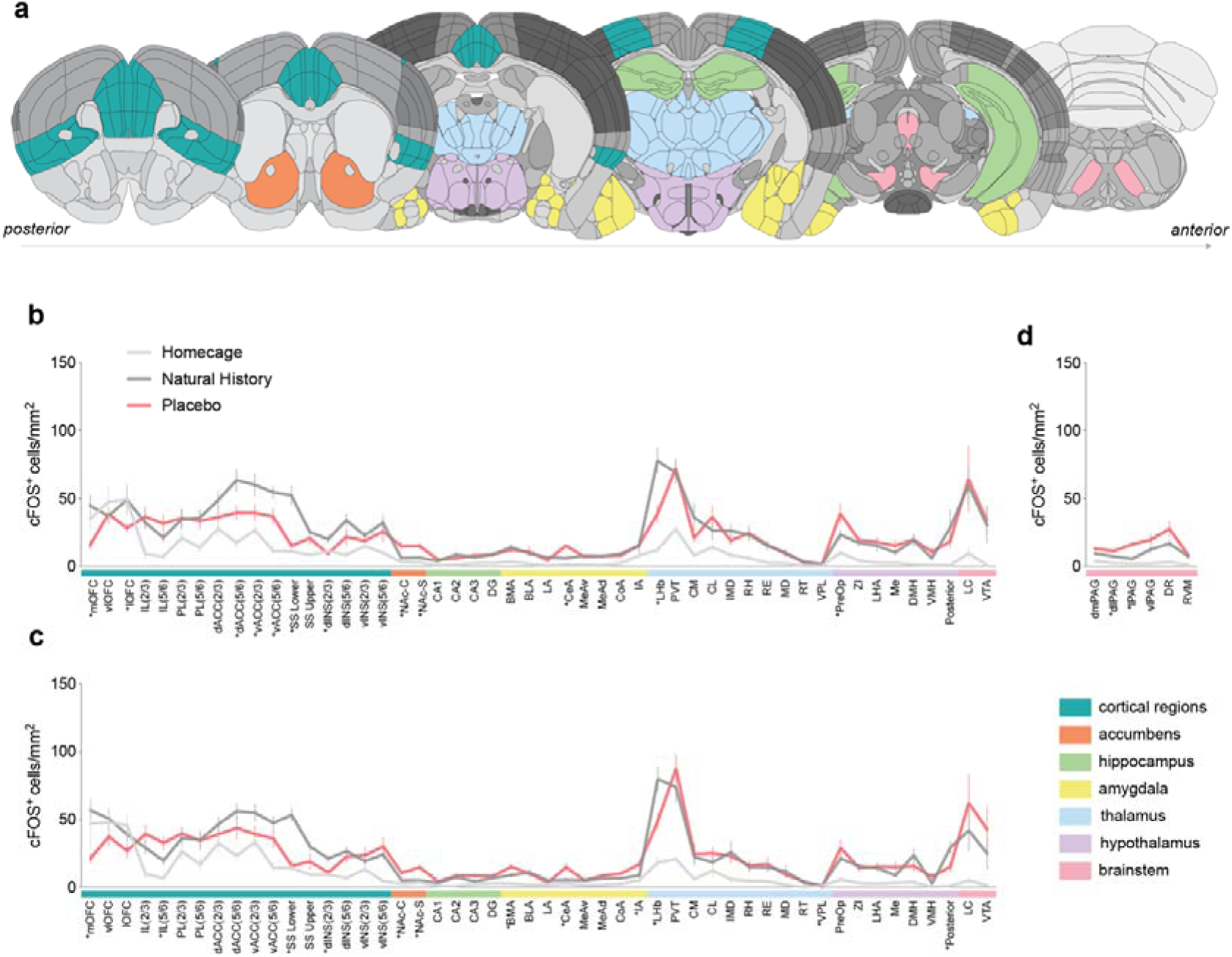
Anatomical distribution and hemispheric patterns of placebo-evoked c-Fos expression. **a** Coronal schematic illustrating the 56 regions of interest (ROIs) in which c-Fos expression was quantified, grouped by major brain regions. **b,c** c-Fos^+^ cell density across ROIs in the left (**b**) and right (**c**) hemispheres following placebo treatment (Mor/Sal, red; n = 10), natural history controls (Sal/Sal, dark grey; n = 10), or home cage controls (light grey; n = 9). **d** c-Fos^+^ cell density across midline brainstem regions. Relative to natural history controls, placebo animals showed decreased c-Fos expression in several cortical regions, including the medial orbitofrontal cortex (left p = 0.0049, g = 1.54; right p = 0.0016, g = 1.82), anterior cingulate cortex (left dACC layer 5/6 p = 0.0254, g = 1.10; left vACC layer 5/6 p = 0.0344, g = 1.03), primary somatosensory cortex lower limb (left p = 0.00045, g = 2.23; right p < 0.0001, g = 3.09), and dorsal insular cortex (left p = 0.0069, g = 1.42; right p = 0.0067, g = 1.38). In contrast, placebo animals exhibited increased c-Fos expression in the lateral periaqueductal gray (p = 0.00023, g = −2.30), nucleus accumbens shell (left p = 0.0057, g = −1.50; right p = 0.00067, g = −2.03), central amygdala (p = 0.00006, g = −2.37), and right basomedial amygdala (p = 0.0400, g = −1.00). ROIs are ordered anatomically from posterior to anterior and identically aligned across hemispheres. Asterisks indicate ROIs that met criteria for both nominal statistical significance (uncorrected p < 0.05) and a large effect size (|g| > 0.8) for the placebo versus natural history comparison. Each point represents group mean ± s.e.m. For a complete list of brain regions and acronyms, see Supplementary Table 1. Full statistical reporting is provided in Supplementary Data 1. Source data are provided as a Source Data file.

In subcortical forebrain structures, the nucleus accumbens core and shell were reduced bilaterally, whereas hippocampal subfields and amygdalar subdivisions largely resembled natural history levels, with only modest, localized differences (**Fig. 2b,c**). Thalamic and hypothalamic nuclei exhibited small, spatially restricted differences without consistent hemispheric differences. In contrast, activity was elevated across midbrain and brainstem regions implicated in descending control, most prominently within the dorsolateral and lateral periaqueductal gray, with additional increases in the dorsal raphe and rostral ventromedial medulla. Increased c-Fos was also observed in locus coeruleus and ventral tegmental area (**Fig. 2d**).

Although naloxone blocked placebo expression behaviorally, broad regional c-Fos activity patterns remained related across conditions. Regional c-Fos densities were moderately associated between placebo and natural history mice (R² = 0.66, slope = 0.65), whereas naloxone-treated placebo mice more closely resembled natural history activity patterns (R² = 0.93, slope = 1.20) (Supplementary Fig. 7a,b). Direct comparison of placebo-associated changes with naloxone-induced shifts nevertheless revealed substantial regional heterogeneity in the direction and magnitude of opioid-dependent modulation (Supplementary Fig. 7c). To quantify these effects, we calculated a naloxone reversal index using hemisphere-combined c-Fos counts, which measured the extent to which naloxone shifted activity toward or away from natural history levels (Supplementary Fig. 7d). Several cortical regions exhibiting placebo-associated suppression, including lower-limb somatosensory cortex, dorsal anterior cingulate cortex layer 5/6, and medial orbitofrontal cortex, showed significant reversal toward natural history levels following naloxone administration (Supplementary Fig. 7e). Similarly, placebo-associated elevations within nucleus accumbens shell and ventrolateral PAG were reduced by naloxone (Supplementary Fig. 7f). In contrast, other regions exhibited little reversal or diverged further from the natural history state following naloxone administration. Paraventricular thalamus and locus coeruleus showed elevated mean c-Fos activity in naloxone-treated mice relative to both placebo and natural history groups, although these differences did not reach statistical significance (Supplementary Fig. 7g). Ventral insular cortex layer 2/3 and ventrolateral orbitofrontal cortex likewise diverged further from natural history levels following naloxone administration (Supplementary Fig. 7h). Together, these findings indicate that opioid receptor blockade selectively alters subsets of placebo-associated neural activity rather than uniformly restoring regional activity to a natural history state.

### Whole-brain functional connectivity analyses identify candidate opioid-sensitive placebo circuits

To determine whether placebo expression altered coordinated activity across regions, we examined whole-brain functional connectivity based on inter-regional correlations in c-Fos density (**Fig. 3a–f**). Functional connectivity matrices revealed marked differences in the large-scale organization of inter-regional correlations between natural history and placebo mice (**Fig. 3a,d**). In natural history mice, correlated activity was broadly distributed across thalamic, hypothalamic, and brainstem regions, with comparatively diffuse organization across cortical areas (**Fig. 3a**). In contrast, placebo mice exhibited increased clustering of positive correlations within cortical and cortical-associated regions, suggesting a redistribution of coordinated activity during placebo expression (**Fig. 3d**). To identify the connections underlying this shift, we visualized the 100 strongest correlations using thresholded network representations, matching network density across conditions (**Fig. 3b,e**). In natural history mice, the strongest connections were dominated by subcortical–subcortical interactions, which comprised 70% of all edges, whereas cortical–cortical interactions accounted for only 8%, with the remaining 22% occurring between cortical and subcortical regions (**Fig. 3b**, Supplementary Table 2). In placebo mice, this organization shifted markedly toward cortical-cortical and corticolimbic connectivity, with cortical–cortical interactions increasing nearly fivefold to 39% and subcortical–subcortical interactions decreasing to 27%, while cortical–subcortical interactions also increased (34%, **Fig. 3e**, Supplementary Table 2). The distribution of connection classes differed significantly between conditions (χ² = 42.08, p < 0.0001), indicating that placebo expression reorganizes coordinated activity away from predominantly subcortical connectivity toward cortical and limbic network integration.

**Figure 3.**
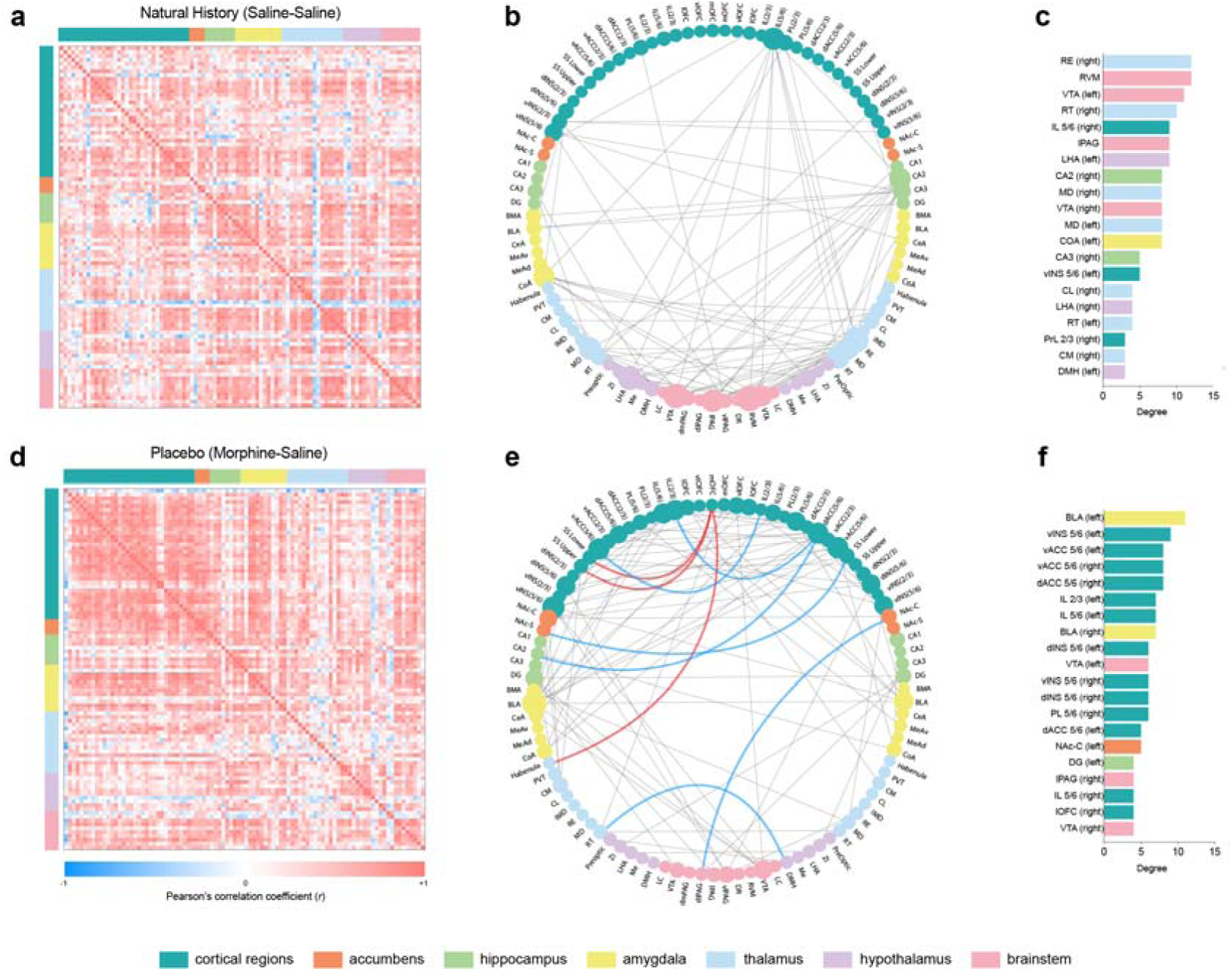
Reorganization of whole-brain functional connectivity during placebo analgesia. a,d. Group-level functional connectivity matrices derived from Pearson correlations of regional c-Fos densities across all quantified brain regions in control (**a**, n = 10) and placebo (d, n = 10) conditions. Each matrix element represents the strength of the correlation between a pair of regions, with warmer colours indicating stronger positive correlations and cooler colours indicating weaker or negative correlations. Regions are ordered and colour-coded by major anatomical division (cortex, accumbens, hippocampus, amygdala, thalamus, hypothalamus, brainstem). **b,e** Circular network representations of functional connectivity for the control (**b**) and placebo (**e**) groups, constructed from the top 100 strongest connections (by absolute Pearson’s r) in each condition. Nodes represent individual brain regions, grouped by anatomical category, and edges represent undirected functional connections. Grey edges indicate connections present within each group’s top-strength network. In the placebo network (**e**), coloured edges highlight connections that differ significantly between groups after multiple-comparison correction (q-value threshold), with red edges indicating increased connectivity and blue edges indicating decreased connectivity relative to control. **c,f** Node degree distributions for the control (**c**) and placebo (**f**) networks, defined as the number of connections per region within the top-100-connection network. Bars are colour-coded by anatomical group, illustrating shifts in network hubs and regional centrality between conditions. Together, these analyses reveal a placebo-associated reconfiguration of functional brain networks, characterized by selective strengthening and weakening of specific inter-regional connections and redistribution of hub regions across cortical and subcortical structures. Source data are provided as a Source Data file.

Given this large-scale redistribution of inter-regional coupling, we next examined whether placebo expression altered the organization of network hubs. Node degree analysis revealed that highly connected regions in natural history mice were dominated by subcortical structures, including RVM, RE, VTA, RT, lPAG, and LHA, together with additional thalamic and limbic regions (**Fig. 3c**). In contrast, placebo mice exhibited a redistribution of highly connected nodes toward cortical and corticolimbic regions, including bilateral basolateral amygdala, ventral insula, ventral anterior cingulate, dorsal anterior cingulate, and infralimbic cortex subdivisions, while midbrain regions such as lPAG and VTA remained integrated within the reorganized network (**Fig. 3f**).

To determine whether opioid receptor blockade restored the placebo-associated network toward a natural history state, we examined whole-brain functional connectivity in naloxone-treated mice (Supplementary Fig. 8a–c). Consistent with our region-specific analysis, across matrices, network topology, and hub organization, the naloxone condition exhibited features distinct from both placebo and natural history states. Despite behavioral blockade of placebo expression, naloxone did not restore the network to a natural history configuration, suggesting that opioid receptor blockade perturbs selective aspects of the placebo-associated network state without globally resetting large-scale functional organization. To further understand how inter-regional coupling changed across network states, we performed signed strength analyses examining edgewise differences between conditions (Supplementary Fig. 9 and Supplementary Data 2). Relative to natural history, placebo expression was associated with widespread reorganization of functional coupling, including strengthened interactions between medial orbitofrontal cortex and habenula, as well as between medial orbitofrontal and somatosensory regions (**Fig. 3e**, red lines; Supplementary Fig. 9a). In contrast, several cortical–subcortical and subcortical interactions were weakened, including dACC layer 2/3–infralimbic cortex layer 2/3, dACC layer 2/3–CA2, lower-limb somatosensory cortex–nucleus accumbens shell, nucleus accumbens core–dlPAG, and dorsomedial hypothalamus–reticular thalamus connections (**Fig. 3e**, blue lines; Supplementary Fig. 9a). Compared with the large-scale edge reorganization observed between placebo and natural history states, naloxone produced comparatively restricted alterations in inter-regional coupling (Supplementary Fig. 9b), while natural history and naloxone networks continued to differ across multiple inter-regional connections (Supplementary Fig. 9c). Together, these findings indicate that placebo expression is associated with large-scale corticolimbic network reorganization, whereas opioid receptor blockade disrupts selective components of this reorganized brain state rather than globally restoring natural history connectivity patterns.

### Excitatory activation of the anterior cingulate cortex blocks placebo analgesia

Although multiple regions differed between placebo and natural history animals, the ACC showed convergent changes across activity, inter-regional coupling, and network centrality. Given the established role of the ACC in expectancy-driven amplification of pain[37, 38], and top-down modulation of nociception[39, 40]. This convergence suggests that functional suppression of cingulate output may contribute to placebo analgesia. We therefore tested whether bidirectional manipulation of ACC activity during the placebo test day would causally influence placebo expression. Mice received bilateral injections of excitatory (hM3Dq), inhibitory (hM4Di), or control virus into the ACC. To control for potential locomotor effects of chemogenetic manipulation, mice were first tested in an open field following CNO administration prior to conditioning. Mice then underwent the standard placebo conditioning procedure, and CNO was administered again on the test day prior to von Frey measurements (**Fig. 4a**). Representative images illustrating bilateral ACC targeting for control, inhibitory, and excitatory groups are shown in **Fig. 4b–e**, with full reconstruction of viral spread across mice provided in Supplementary Fig. 10.

**Figure 4.**
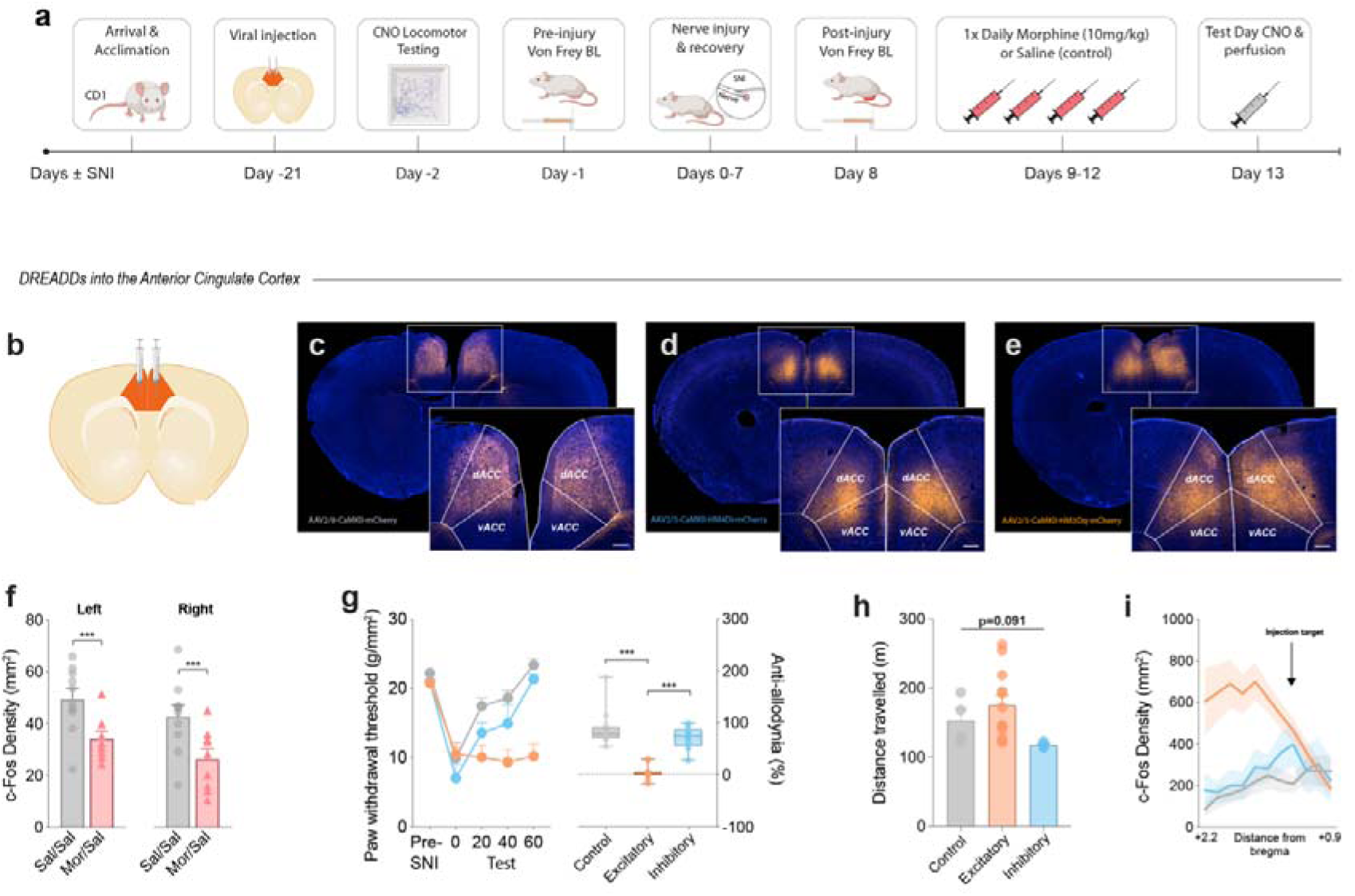
Excitatory DREADD-mediated activation of the anterior cingulate cortex blocks placebo analgesia. **a** Experimental timeline. CD1 mice received bilateral stereotaxic injections of control, excitatory (hM3Dq), or inhibitory (hM4Di) DREADDs fused to mCherry. Three weeks later, mice underwent the standard placebo conditioning protocol. On the test day, clozapine-N-oxide (CNO; i.p.) was administered immediately prior to von Frey testing, and mice were perfused 90 min later for whole-brain c-Fos analysis. **b–e** Schematic and representative images illustrating bilateral stereotaxic injections targeting the anterior cingulate cortex (ACC), showing mCherry-labelled control (**c**), inhibitory (**d**), and excitatory (**e**) DREADD expression. Insets show magnified views of the injection site. Scale bars in insets represent 200 μm. **f** Quantification of c-Fos-positive cell density within the ACC from the whole-brain analysis, shown separately for left and right hemispheres. Placebo (Mor/Sal, n = 9) mice exhibited reduced ACC c-Fos expression relative to natural history controls (Sal/Sal, n = 9) in both hemispheres (left, t = 3.09, d.f. = 15.56, p = 0.0072; right, t = 2.78, d.f. = 16.93, p = 0.013). **g** Paw withdrawal thresholds to von Frey stimulation measured at pre-injury baseline (Pre-SNI) and at 0, 20, 40, and 60 min following CNO administration on the test day. Withdrawal thresholds differed across groups over time (group × time interaction, F(8,120) = 10.01, p < 0.0001). Right panel shows percentage anti- allodynia relative to baseline. Anti-allodynia differed between groups (Welch’s ANOVA, F(2,16.45) = 53.01, p < 0.0001), with the excitatory group (n = 12) differing from both control (n = 12, p < 0.0001) and inhibitory (n = 9, p < 0.0001) groups, whereas control and inhibitory groups did not differ (p = 0.4205). **h** Locomotor activity measured following CNO administration. No significant differences were observed between groups (one-way ANOVA, F(2,17) = 2.77, p = 0.091). Control, n = 4 mice; Excitatory, n = 8 mice; Inhibitory, n = 4 mice. **i** Spatial distribution of c-Fos density relative to bregma following chemogenetic manipulation, demonstrating localized modulation of neuronal activity near the ACC injection site. c-Fos density varied as a function of experimental group (n = 8 mice/group) and distance from bregma (group × bregma interaction, F(20,194) = 4.04, p < 0.0001). Shaded areas represent s.e.m. Full statistical reporting is provided in Supplementary Data 1. Source data are provided as a Source Data file.

Given that the ACC showed reduced c-Fos expression in the placebo group (**Fig. 4f**), we hypothesized that increasing ACC activity would block placebo behavior. Control and inhibitory (hM4Di) groups displayed robust placebo analgesia, with increased withdrawal thresholds following CNO administration on the test day. In contrast, excitatory ACC activation abolished placebo expression, with thresholds remaining near post-injury levels across the testing period (**Fig. 4g**). Importantly, chemogenetic manipulations did not alter mechanical withdrawal thresholds in the absence of placebo conditioning, indicating that the observed effects were specific to placebo expression rather than nonspecific modulation of mechanical sensitivity (Supplementary Fig. 11). Effects were not explained by nonspecific changes in locomotor activity, as open field testing conducted prior to conditioning showed no significant differences in distance travelled between groups following CNO administration, despite a modest, non-significant increase in the excitatory group (**Fig. 4h**). Spatial analysis of c-Fos along the anterior–posterior axis confirmed that modulation was centered near the ACC injection site, with divergent profiles for excitatory and inhibitory manipulations (**Fig. 4i**, Supplementary Fig. 12). Together, this shows that increasing ACC activity during the placebo test abolishes placebo analgesia, whereas inhibitory manipulation does not further enhance the response, indicating that reduced ACC activity permits, rather than rate-limits, placebo expression.

### μ-opioid receptor–expressing neurons in the ACC are necessary for placebo analgesia expression

Given that placebo expression was μ-opioid-dependent and ACC activation abolished the response, we next tested whether μ-opioid receptor–expressing neurons within the ACC mediate placebo expression. Thus, we selectively ablated MOR+ cells using dermorphin–saporin (Derm-Sap), a conjugated toxin that is internalized by MOR-expressing neurons and induces cell death following receptor-mediated uptake[41]. Bilateral ACC administration of Derm-Sap, compared with blank saporin (Blank-Sap) lacking the dermorphin targeting peptide, resulted in a reduction of MOR protein levels within ACC tissue, confirming selective depletion of MOR-expressing neurons (**Fig. 5a,b**, Supplementary Fig. 13). Morphine conditioning produced comparable anti-allodynia across conditioning days in Blank-Sap and Derm-Sap groups, indicating that ablation of MOR-expressing ACC neurons did not impair acute morphine analgesia (**Fig. 5c**). However, during placebo testing, Blank-Sap mice exhibited robust recovery of withdrawal thresholds, whereas Derm-Sap–treated animals failed to express placebo analgesia (**Fig. 5d**), suggesting that μ-opioid receptor–expressing neurons in the ACC are required for placebo analgesia.

**Figure 5.**
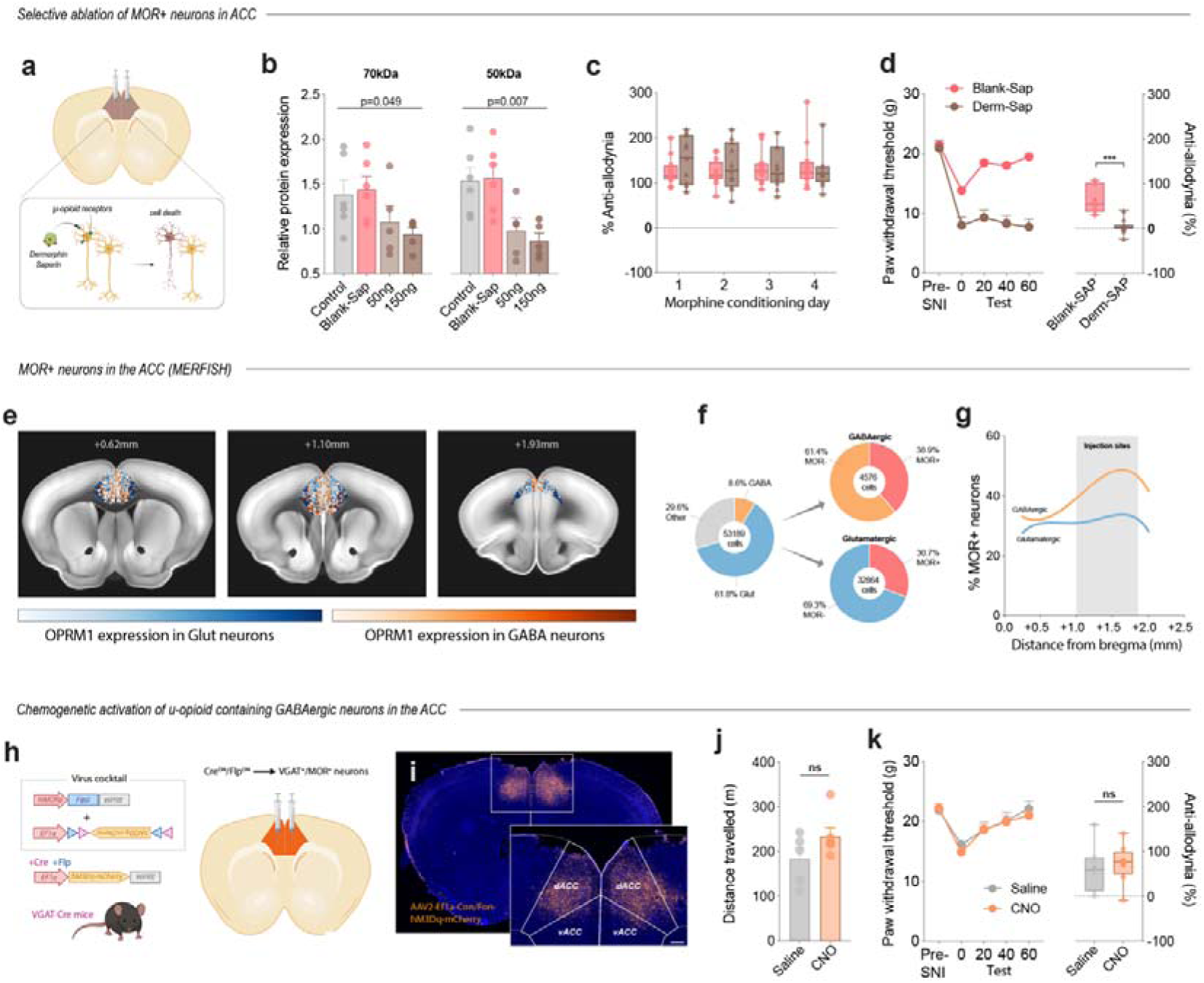
Ablation of μ-opioid receptor–expressing neurons in the anterior cingulate cortex prevents placebo analgesia in morphine conditioned mice. Experimental strategy for selective ablation of μ-opioid receptor (MOR; OPRM1)-expressing neurons in the anterior cingulate cortex (ACC) using dermorphin-saporin (Derm-Sap). Control animals received blank saporin (Blank-Sap). **b** Western blot quantification of MOR protein levels (70 kDa and 50 kDa bands) in ACC tissue following Blank-Sap or two concentrations of Derm-Sap. MOR protein levels differed across groups for both the 70 kDa (Welch’s ANOVA, F(3,9.37) = 3.83, p = 0.049) and 50 kDa bands (Welch’s ANOVA, F(3,9.76) = 7.38, p = 0.0071). Control, n = 6 mice; Blank-Sap, n = 6 mice; 50 ng Derm-Sap, n = 5 mice; 150 ng Derm-Sap, n = 5 mice. **c** Percentage anti-allodynia across morphine conditioning days following selective MOR^+^ neuron ablation. MOR^+^ neuron ablation did not alter the acquisition of conditioned analgesia during training (group × time interaction, F(3,62) = 1.53, p = 0.21). Blank-Sap, n = 12 mice; Derm-Sap, n = 11 mice. **d** Paw withdrawal thresholds (left) and anti-allodynia (right) during placebo testing in Blank-Sap (n = 12 mice) and Derm-Sap (n = 12 mice) groups. Withdrawal thresholds differed between groups over time (group × time interaction, F(4,76) = 15.98, p < 0.0001), and Derm-Sap treatment reduced placebo expression relative to Blank-Sap controls (t = 4.36, d.f. = 6.37, p = 0.0041). **e-g** Characterization of *Oprm1* expression in ACC neurons using a publicly available single-mouse MERFISH dataset. **e** Representative coronal sections illustrating Oprm1-expressing cells across the ACC at multiple bregma levels. **f** Quantification of *Oprm1* expression within glutamatergic and GABAergic neuronal populations in the ACC, expressed as percentages and absolute counts. **g** Spatial distribution of *Oprm1^+^*neurons along the anterior-posterior axis. **h** Chemogenetic strategy targeting *Oprm1*-expressing GABAergic neurons in the ACC using hM3Dq DREADDs. **i** Representative fluorescence images showing hM3Dq-mCherry expression within the ACC. Scale bar in inset represents 200 μm. **j** Chemogenetic activation of ACC VGAT/*Oprm1* neurons did not significantly alter locomotor activity (t = 1.59, d.f. = 7.89, p = 0.152). Saline, n = 5 mice; CN), n = 6 mice **k,l** Paw withdrawal thresholds during placebo testing (**k**) and percentage anti-allodynia (**l**) following saline (n = 7 mice) or CNO (n = 8 mice) administration. Chemogenetic activation of ACC VGAT/*Oprm1* neurons did not alter placebo expression (group × time interaction, F(5,13) = 0.38, p = 0.824; anti-allodynia, t = 0.32, d.f. = 11.67, p = 0.756). Data are presented as mean ± s.e.m. Individual data points are shown. *p < 0.05, **p < 0.01, ***p < 0.001, ****p < 0.0001. Full statistical reporting is provided in Supplementary Data 1. Source data are provided as a Source Data file.

To further characterize MOR-containing neurons in the ACC, we analyzed a publicly available MERFISH data set from the Allen Brain Cell Atlas[42]. *Oprm1* was distributed across both glutamatergic and GABAergic neuronal classes, with labeled cells present across dorsal and ventral ACC at multiple rostral-caudal levels (**Fig. 5e**). Across all ACC neurons, 61.8% were glutamatergic and 8.6% were GABAergic, with the remainder classified as other cell types. Within these populations, *Oprm1* was detected in ∼30.7% of glutamatergic neurons and ∼38.9% of GABAergic neurons (**Fig. 5f**). Mapping along the anterior–posterior axis showed that MOR-positive neurons were broadly distributed across the ACC, with both glutamatergic and GABAergic populations represented across the sampled levels (**Fig. 5g**). This pattern was consistent with our RNAscope validation in ACC tissue, which also detected *Oprm1* transcripts in both glutamatergic and GABAergic neurons, with a higher proportion of GABAergic neurons expressing *Oprm1* (Supplementary Fig. 14).

Although GABAergic neurons comprised a smaller fraction of ACC cells overall, a comparable proportion of inhibitory and excitatory neurons expressed *Oprm1*, suggesting that μ-opioid signaling might preferentially recruit local inhibitory microcircuits. Given that activation of ACC excitatory neurons disrupted placebo expression, we next asked whether perturbing opioid-sensitive inhibitory neurons within the ACC was sufficient to alter conditioned analgesia. To test this, we selectively targeted MOR-expressing GABAergic neurons using an intersectional Cre/Flp strategy in VGAT-Cre mice and expressed an hM3Dq DREADDs within this population (**Fig. 5h,i**). Chemogenetic activation of VGAT*^Oprm1^* neurons did not alter locomotor activity following CNO administration (**Fig. 5j**), arguing against nonspecific behavioral effects. Paw withdrawal thresholds increased similarly following saline and CNO administration, and percent anti-allodynia during placebo testing did not differ between conditions (**Fig. 5k**). Thus, selective activation of μ-opioid receptor–expressing GABAergic neurons in the ACC was not sufficient to enhance placebo analgesia, indicating that μ-opioid signaling within the ACC likely engages a broader neuronal population rather than a single inhibitory subset.

### Chemogenetic manipulation of the basomedial amygdala and paraventricular thalamus during placebo expression

Having identified the ACC as a critical node, we next examined whether downstream limbic and thalamic structures contribute to placebo expression. The basomedial amygdala (BMA) and paraventricular thalamus (PVT) integrate affective and contextual information and are well positioned to influence pain-related behavior through interactions with cortical, thalamic, and brainstem circuits[43–47]. Given the established μ-opioid regulation of midline thalamic–amygdala circuits [48], and the emergence of the PVT and BMA as candidate opioid-sensitive nodes in our c-Fos and network analyses (Supplementary Figs. 7 and 9), we tested their contribution to placebo expression using chemogenetic perturbation.

Mice received bilateral injections of control, excitatory (hM3Dq), or inhibitory (hM4Di) DREADDs into the BMA (**Fig. 6a–d**). Whole-brain activity mapping identified a localized increase in c-Fos expression within the right BMA during placebo expression (**Fig. 6e**), raising the possibility that activity within this region contributes to placebo analgesia. We therefore tested whether bidirectional manipulation of BMA activity alters placebo behavior. Paw withdrawal thresholds recovered similarly across control, excitatory, and inhibitory groups following CNO administration, and percent anti-allodynia did not differ between conditions (**Fig. 6f**). Locomotor activity did not differ between groups following CNO administration (**Fig. 6g**). CNO bidirectionally modulated BMA activity, with increased c-Fos in the excitatory group and reduced labeling in the inhibitory group, centered at the injection site (**Fig. 6h**, Supplementary Fig. 12).

**Figure 6.**
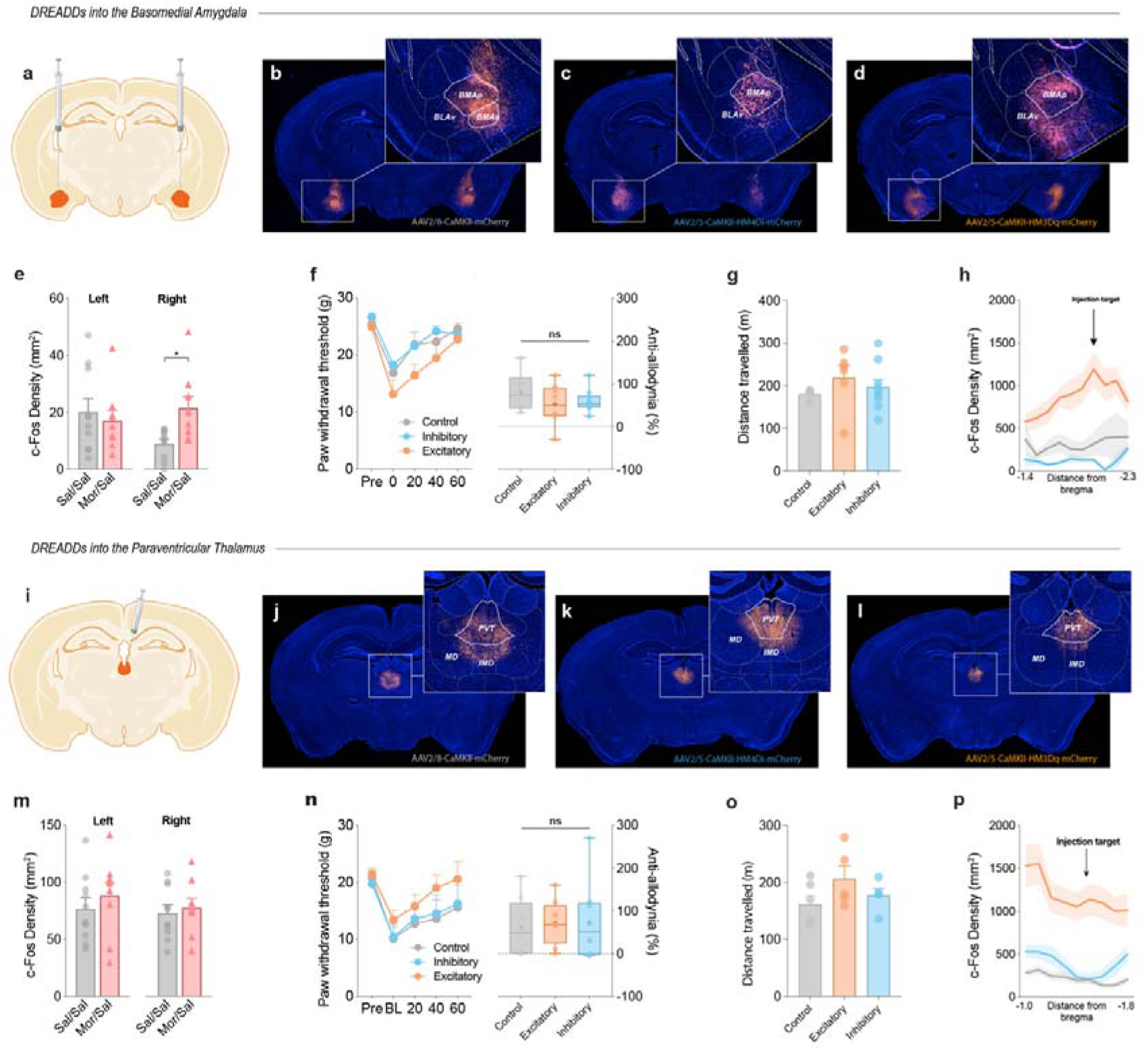
Chemogenetic modulation of the basomedial amygdala and paraventricular thalamus does not alter placebo analgesia. **a** Schematic illustrating bilateral stereotaxic injection of control, excitatory (hM3Dq), or inhibitory (hM4Di) DREADDs into the basomedial amygdala (BMA). **b–d** Representative images showing mCherry-labelled control (**b**), inhibitory (**c**), and excitatory (**d**) DREADD expression within the BMA. Insets show magnified views of the injection site. Scale bars in insets represent 200 μm. **e** Quantification of c-Fos-positive cell density within the BMA from the whole-brain analysis, shown separately for left and right hemispheres. Placebo animals (Mor/Sal, n = 9 mice) exhibited increased c-Fos expression within the right BMA relative to natural history controls (Sal/Sal, n = 10 mice; t = 2.91, d.f. = 10.6, p = 0.01), whereas no difference was observed in the left hemisphere (t = 0.54, d.f. = 16.48, p = 0.60). **f** Paw withdrawal thresholds to von Frey stimulation measured at pre-injury baseline and at 0, 20, 40, and 60 min following CNO administration on the test day. Withdrawal thresholds did not differ between groups over time (group × time interaction, F(8,88) = 0.93, p = 0.494). Right panel shows percentage anti-allodynia relative to baseline. Anti-allodynia did not differ between groups (Welch’s ANOVA, F(2,21) = 0.90, p = 0.42). Control, n = 7 mice; Excitatory, n = 8 mice; Inhibitory, n = 10 mice. **g** Locomotor activity measured following CNO administration. No significant differences were observed between groups (one-way ANOVA, F(2,16) = 0.65, p = 0.54). Control, n = 9 mice; Excitatory, n = 4 mice; Inhibitory, n = 6 mice. **h** Spatial distribution of c-Fos density relative to bregma following chemogenetic manipulation, demonstrating bidirectional modulation of neuronal activity centred on the BMA injection site. c-Fos density differed between experimental groups (F(2,20) = 35.14, p < 0.0001), with no group × bregma interaction (F(8,122) = 1.53, p = 0.09). Shaded areas represent s.e.m. Control, n = 7 mice; Excitatory, n = 8 mice; Inhibitory, n = 8 mice. **i** Schematic illustrating bilateral stereotaxic injections of control, excitatory (hM3Dq), or inhibitory (hM4Di) DREADDs into the paraventricular thalamus (PVT). **j–l** Representative images showing mCherry-labelled control (**j**), inhibitory (**k**), and excitatory (**l**) DREADD expression within the PVT. Insets show magnified views of the injection site. Scale bars in insets represent 200 μm. **m** Quantification of c-Fos-positive cell density within the PVT from the whole-brain analysis, shown separately for left and right hemispheres. No differences in PVT c-Fos expression were observed between placebo (Mor/Sal, n = 9 mice) and natural history (Sal/Sal, n = 10 mice) groups (left, t = 0.78, d.f. = 15.93, p = 0.44; right, t = 0.45, d.f. = 16.77, p = 0.66). **n** Paw withdrawal thresholds measured as above. Withdrawal thresholds did not differ between groups over time (group × time interaction, F(8,84) = 0.48, p = 0.87). Right panel shows percentage anti-allodynia relative to baseline. Anti-allodynia did not differ between groups (Welch’s ANOVA, F(2,21) = 0.07, p = 0.94). Sample size, n = 8 mice/group. **o** Locomotor activity measured following CNO administration. No significant differences were observed between groups (one-way ANOVA, F(2,13) = 1.75, p = 0.21). Control, n = 5 mice; Excitatory, n = 6 mice; Inhibitory, n = 5 mice. **p** Spatial distribution of c-Fos density relative to bregma following chemogenetic manipulation, demonstrating bidirectional modulation of neuronal activity centred on the PVT injection site. c-Fos density varied as a function of experimental group and distance from bregma (group × bregma interaction, F(8,165) = 1.86, p = 0.028). Sample size, n = 8 mice/group. Shaded areas represent s.e.m. Data are presented as mean ± s.e.m. Individual data points are shown. *p < 0.05, **p < 0.01, ***p < 0.001, ****p < 0.0001. Full statistical reporting is provided in Supplementary Data 1. Source data are provided as a Source Data file.

We next tested involvement of the paraventricular thalamus, where mice received bilateral injections of control, excitatory (hM3Dq), or inhibitory (hM4Di) DREADDs into the PVT prior to placebo conditioning (**Fig. 6i–l**). Whole brain c-Fos quantification revealed comparable levels of PVT activity across placebo and natural history groups during placebo expression (**Fig. 6m**). Nevertheless, prior work implicating midline thalamic circuits in placebo[4] and expectation-dependent analgesia[49], prompted causal testing of PVT activity. Paw withdrawal thresholds recovered similarly across control, excitatory, and inhibitory groups following CNO administration, and percent anti-allodynia did not differ between conditions (**Fig. 6n**). Locomotor activity also remained comparable between groups, arguing against nonspecific behavioral effects of PVT manipulation (**Fig. 6o**). Mapping of c-Fos density along the anterior–posterior axis showed peak labeling at the injection coordinates, consistent with localized modulation of PVT activity (**Fig. 6p**, Supplementary Fig. 12). Together, these results indicate that, despite their anatomical positioning and involvement in affective and expectation-related processing, these subcortical relays are not required for placebo analgesia, further narrowing the circuit to cortical control mechanisms.

## Discussion

Placebo analgesia engages endogenous opioid signaling and cortical circuits in humans[11, 50], but how these mechanisms are expressed across brain-wide networks in chronic pain states remains unclear. Recent rodent studies have begun to identify the circuits that support conditioned placebo-like analgesia but have largely relied on targeted manipulations of candidate pathways, leaving unresolved how placebo responses are organized at the whole-brain level, particularly in chronic pain. Here we show that robust, context-dependent placebo analgesia can be established in a chronic neuropathic pain state in mice and that its expression reflects a coordinated reorganization of brain-wide activity rather than a simple reduction in nociceptive drive. Across conditions, placebo expression was associated with a redistribution of activity, characterized by the suppression of cortical regions involved in sensory and interoceptive processing and by increased engagement of midbrain and brainstem structures implicated in descending pain modulation. This pattern is consistent with models in which placebo reduces ascending nociceptive signaling while enhancing descending control[8, 11] but extends these models by showing that placebo expression is associated with selective redistribution of activity and inter-regional coupling across cortical, corticolimbic, and descending modulatory circuits rather than isolated changes within individual regions. This interpretation is further supported by the observation that placebo did not produce a uniform scaling of activity across the brain. Instead, deviations from the natural history state were restricted to a subset of regions, indicating a redistribution of activity rather than a global shift.

Network reconfiguration was characterized by a shift away from subcortically dominated connectivity toward increased cortical and corticolimbic coupling. In natural history animals, the strongest connections were largely confined to subcortical circuits, whereas placebo expression increased intracortical and corticolimbic interactions and reduced subcortical coupling. These findings align with human imaging work showing that placebo engages distributed cortical networks, including prefrontal and cingulate regions, and alters connectivity with brainstem pain-modulatory systems[8, 51], but extend this view by showing how these changes emerge from a redistribution of network architecture rather than uniform modulation of connectivity strength. Importantly, the network shift that we observe was not driven by widespread changes in node centrality. Node-level effects were sparse after correction for multiple comparisons, indicating that placebo was not associated with global reassignment of hub structure. Instead, changes in network organization were supported by selective reweighting of inter-regional coupling, as revealed by signed strength analyses. This suggests that placebo expression is associated with targeted differences in connectivity between specific cortical and subcortical nodes rather than large-scale reorganization of network hubs. Notably, this unbiased, brain-wide approach did not identify a novel subcortical driver of placebo analgesia, but instead converged on cortical regions, particularly the ACC, as dominant contributors to network reorganization.

The ACC was the only region showing convergent changes across activity, inter-regional coupling, and network centrality, and causal manipulations confirmed that increasing ACC activity abolished placebo analgesia. This is consistent with a large body of human work implicating the ACC in expectancy-driven modulation of pain and placebo responses[8, 11, 52], as well as rodent studies linking ACC activity to top-down control of nociception[53, 54]. Our findings extend these observations by showing that reduced ACC activity is permissive for placebo expression in chronic pain, rather than a downstream relay within a distributed circuit. Importantly, the ACC emerged as the central node in an unbiased, whole-brain analysis, reinforcing its role as a primary cortical locus for placebo expression rather than simply a component of a distributed circuit. Systemic and local manipulations demonstrated that placebo expression depends on μ-opioid receptor activity and that ablation of MOR-expressing neurons in the ACC abolishes the response. These findings are consistent with human PET studies showing endogenous opioid release during placebo analgesia[10, 11, 50] and suggest that μ-opioid signaling within cortical circuits is necessary for translating learned expectations into behavioral analgesia. However, selective activation of MOR-expressing GABAergic neurons was insufficient to block placebo responses, suggesting that μ-opioid signaling likely engages a broader cellular ensemble rather than a single inhibitory pathway. Together, our findings identify the ACC as a primary cortical gate for placebo expression in chronic pain, rather than a downstream relay within a distributed circuit.

Direct cross-species comparison of placebo analgesia and nocebo hyperalgesia in humans and rats demonstrates that placebo responses exhibit more conserved circuitry across species than nocebo responses, with overlapping placebo-related connectivity concentrated in the ACC, amygdala, and nucleus accumbens, and hub regions including the ACC and PAG-related pain-modulatory structures[49]. Consistent with this, rodent studies have identified projection-defined cortical circuits that can drive placebo analgesia. For instance, μ-opioid signaling in the mPFC engages placebo responses through an mPFC→vlPAG pathway in pharmacologically conditioned neuropathic rats[13]. In contrast, studies in pain-naive animals show that an ACC→pontine projection is required for expectation-driven placebo responses[12], and more recent work demonstrates that cortical input from the ACC and mPFC to the vlPAG is necessary for placebo analgesia, with endogenous opioid signaling in the vlPAG mediating its expression. Notably, these studies rely on projection-specific manipulations of ACC outputs to establish that distinct cortical pathways can recruit descending analgesic circuits, but do not test whether ACC activity itself is required for placebo expression. Our study extends this growing literature in three ways. First, rather than beginning with a candidate descending pathway, we used an unbiased whole-brain activity and network analysis and still converged on the ACC. Second, we show that increasing ACC activity is sufficient to abolish placebo expression, indicating that ACC activity gates the behavioral response. Third, selective ablation of ACC MOR^+^ neurons eliminates placebo analgesia without disrupting acute morphine analgesia, identifying a cortical μ-opioid-dependent population required for placebo expression. Thus, our data complement projection-defined mPFC→vlPAG work by identifying the upstream cortical gate through which placebo analgesia is organized in chronic neuropathic pain.

The necessity of the ACC for placebo in our model also aligns with evidence that expectation can drive pain in opposing directions through distinct cortical mechanisms. Both placebo and nocebo states ultimately engage PAG-dependent descending control, yet our prior work showed that cholecystokinin (CCK) signaling within ACC to lateral PAG (lPAG) circuits promotes nocebo hypersensitivity[55], whereas the present study identifies ACC μ-opioid signaling as necessary for placebo analgesia. This is consistent with behavioral pharmacology studies in humans showing that CCK facilitates nocebo hyperalgesia, whereas endogenous opioid signaling mediates placebo analgesia, reflecting opposing neurochemical regulation over these states[7]. Whether these signals arise from distinct neuronal populations or differential recruitment of shared cells remains unresolved. One interpretation is that these opposing states are routed through distinct ACC output pathways that converge onto common brainstem targets, with placebo engaging inhibitory control pathways and nocebo engaging facilitatory pathways. Under this model, broad chemogenetic activation of ACC excitatory neurons may recruit pro-nociceptive nocebo-related circuits, including the previously identified ACC→lPAG CCKergic pathway, thereby overriding placebo-associated analgesic signaling. While the present data do not resolve these specific projections, they identify the ACC as a cortical hub through which these opposing modulatory states may be organized before converging onto shared descending circuitry.

Although placebo expression altered activity across multiple brain regions, causal manipulations of subcortical nodes did not reveal a comparable role for these structures. Neither the basomedial amygdala nor the paraventricular thalamus was required for placebo expression, despite their anatomical positioning and known involvement in affective and expectation-related processes[43, 44, 46, 47]. This suggests that while these regions may participate in shaping network dynamics, they are not essential drivers of placebo analgesia in this context, further narrowing the circuit to cortical control mechanisms. Naloxone abolished placebo behavior but did not revert brain-wide activity or network organization to the natural history state. Instead, its effects were restricted to a subset of regions and connections, indicating that opioid signaling does not globally define the placebo network but selectively modulates specific components of an already reorganized system. This dissociation between behavioral reversal and partial network perturbation suggests that once established, the placebo state reflects a distributed configuration that is not easily undone by blocking a single neuromodulatory pathway.

Together, our findings do not fit a simple gain-like model of placebo analgesia in chronic pain. Placebo did not produce a uniform suppression of pain-related activity but was instead associated with a selective redistribution of network engagement, with reduced activity in cortical regions linked to sensory and interoceptive processing and increased coordination across cortical and corticolimbic circuits, alongside recruitment of descending control systems. These changes were not accompanied by widespread shifts in node centrality, indicating that placebo does not reflect a global reweighting of the network, but rather targeted adjustments in coupling between specific regions. Across these analyses, the ACC consistently emerged as the dominant node, and causal manipulations show that its activity gates placebo expression through μ-opioid signaling. This places the ACC as an upstream control point through which learned expectations are translated into behavioral analgesia in the context of chronic pain. Placebo analgesia, therefore, reflects an active reorganization of brain-wide interactions embedded within chronic pain circuitry, rather than a passive reduction in nociceptive drive, and provides a systems-level account for how cognitive context can override established pain states.

## Methods

### Animals and housing

Most experiments were performed in naïve, adult male CD-1 (ICR:Crl) (6–12 weeks of age) obtained from Charles River Laboratories (Saint-Constant, QC, Canada). In a subset of experiments, female CD-1 mice and male C57BL/6NCrl mice were included to assess generalizability across sex and strain. In experiments requiring cell-type–specific manipulations, VGAT-Cre mice (Slc32a1-IRES-Cre; Jackson Laboratory, stock #016962) were used to target GABAergic neuronal populations. VGAT-Cre mice were maintained on a C57BL/6 background, bred in-house, and age-matched to wild-type cohorts. Mice were group-housed (n = 2 – 4 per cage) on individually ventilated cage racks (Techniplast North America) under a 12:12 h light/dark cycle (lights on at 07:00 h) in a temperature-controlled (22 ± 2°C) room with access to food (2019 Teklad) and water *ad libitum*. Mice were acclimated to the vivarium for at least one week prior to experimentation, and all mice were handled daily for 5 minutes each for at least 3 days prior to commencement of baseline testing to reduce stress. Mice were randomly assigned to experimental groups, and all behavioral testing and analysis were conducted by experimenters blinded to treatment condition. Unless otherwise stated, experiments were performed across at least three independent cohorts, with balanced representation of experimental and control groups within each cohort to control for batch effects. All procedures adhered to Canadian Council on Animal Care guidelines and were approved by the University of Toronto Animal Care Committee.

### Mechanical sensitivity (von Frey) testing

Prior to the spared nerve injury (SNI), baseline mechanical measurements were taken from the left and right hind paws. The mice then underwent the SNI procedure on their left leg and were given 7 days to recover. Mechanical sensitivity was assessed using von Frey filaments (Stoelting Touch Test Sensory Evaluator Kit no. 2 to no. 9; ranging from ∼0.015 g to ∼1.3 g of force) and the Simplified Up and Down (SUDO) method to estimate 50% paw withdrawal thresholds in pressure units (in grams per square millimeter)[56]. Mice were placed in Plexiglas cubicles on an elevated wire mesh platform and allowed to habituate undisturbed for 60 minutes. Filaments were applied to the plantar surface of the hind paw until bending and held for up to 3 seconds. A positive response was defined as rapid withdrawal, flinching, or licking of the paw. Measurements were obtained from both hind paws and averaged across two trials per paw. Testing was conducted during the light phase (08:00 h –12:00 h) under consistent conditions by the same experimenter within each cohort. Mechanical sensitivity was analyzed as raw withdrawal thresholds, which served as the primary outcome measure. For comparisons across animals and cohorts, responses were also expressed as percent anti-allodynia relative to baseline (see below for calculation method), which yielded consistent results across conditions.

### Spared Nerve Injury

Neuropathic hypersensitivity was induced using spared nerve injury (SNI), a well-established model of chronic neuropathic pain[57]. Mice were anesthetized with isoflurane (4% induction, 1.5–2% maintenance in oxygen), and toe pinch reflexes were assessed before and during the procedure. Following a skin incision of the left hind limb, blunt dissection of the lateral thigh exposed the three terminal branches of the sciatic nerve. The common peroneal and tibial branches were firmly ligated with 6-0 silk sutures (Ethicon, Somerville, NJ) and then transected distal to the ligature; approximately 2 mm of each distal stump was excised while leaving the sural branch intact. All overlying muscle and skin layers were closed separately using 6-0 coated vicryl sutures (Ethicon). No analgesics were administered pre- or post-operatively to prevent disruption of the development of the pain state. Mice were allowed to recover in their home cages for 7 days following surgery.

### Pharmacological conditioning paradigm

Placebo analgesia was induced using a pharmacological conditioning procedure following SNI. After recovery, mice underwent four consecutive days of conditioning (days 7–10 post-injury), during which contextual (testing environment), tactile (intraperitoneal injection), and procedural cues were paired with an analgesic drug. On each conditioning day, baseline mechanical thresholds were measured, followed by intraperitoneal injection of morphine sulfate (10 mg/kg, i.p.; CDMV, Quebec). Mechanical sensitivity was reassessed at 20-, 40-, and 60- minutes post-injection to confirm drug-induced analgesia. Control (i.e., Natural History) mice received saline (0.9%) in place of morphine on days 7–10 post-injury. On the test day (day 11), mice were returned to the same context and underwent the same procedure as conditioning, except that morphine was replaced with either saline or an opioid receptor antagonist. Naloxone hydrochloride (1 mg/kg, i.p.; Sigma Aldrich Canada, Oakville, ON) was used as a non-selective opioid receptor antagonist. Pharmacological specificity was assessed using receptor-selective antagonists, including CTOP (μ-opioid receptor antagonist; 1 mg/kg), nor-BNI (κ-opioid receptor, 5 mg/kg), and naltrindole (δ-opioid receptor, 5 mg/kg). Antagonists were administered intraperitoneally 15 min prior to behavioral testing to ensure peak receptor occupancy at the time of assessment. Doses were selected based on prior literature demonstrating receptor-selective blockade[58]. Experimental groups consisted of placebo (morphine conditioning, saline test), antagonist (morphine conditioning, antagonist test), and natural history (saline conditioning, saline test). In separate experiments, alternative conditioning agents were used, including a lower morphine dose (5 mg/kg, i.p.) and gabapentin (40 mg/kg or 100 mg i.p.), to test whether conditioned responses generalized across dose and pharmacological classes.

### Dose-extension paradigm

To assess how dosing regimen influences analgesic responses across sessions, mice received repeated morphine administrations under either continuous or intermittent schedules. Mechanical thresholds were measured each day immediately prior to injection and at the same post-injection timepoints described above using the von Frey assay. In the continuous group, morphine (10 mg/kg, i.p.) was administered daily through D9. In the intermittent group, morphine was administered on alternating days, with saline substituted on D5, D7, and D9.

### Hot plate testing

To assess thermal nociceptive thresholds during pharmacological conditioning, a hot-plate assay was conducted at 55°C. Baseline latency was recorded prior to the initiation of pharmacological conditioning. Mice were habituated for 30 minutes before injection and tested 30 minutes post-injection. Upon placement on the heated surface, latency (in seconds) to the first nocifensive response (e.g., hind paw fluttering, licking, or withdrawal) was recorded, after which mice were immediately removed. Four of five paradigms consisted of four consecutive days of morphine conditioning (days 1–4), followed by a test day (day 5) during which saline or naloxone was administered to assess conditioned responding. In the initial paradigm, mice were injected and returned to their home cage without contextual cues prior to testing. A natural history control group underwent hot plate testing for five consecutive days without injections or cues. Subsequent paradigms incorporated contextual conditioning to strengthen associative learning. Mice were placed in designated cubicles pre- and post-injection before hot-plate testing. Visual and gustatory cues were later added to reinforce conditioning, and in the final iteration, conditioning was extended to six days with a preferred palatable reward used as an additional contextual cue. Across all paradigms, thermal sensitivity was consistently measured 30 minutes post-injection.

### Open field locomotor assessment

To determine whether DREADD-mediated activation or inhibition altered baseline locomotor behavior, mice underwent open field testing approximately three weeks following viral injection and two days prior to nerve injury. Mice received clozapine-n-oxide (CNO; 3 mg/kg, i.p.) and were placed immediately into an open field arena (40 × 40 × 40 cm), where locomotor activity was recorded from above for 60 min under low-light conditions. Total distance traveled was quantified using DeepLabCut (v3.0) and Simple Behavioral Analysis (SIMBA, v1.55), tracking the position of the base of the tail. Locomotor activity was compared across control, hM4Di, and hM3Dq groups.

### Tissue collection and perfusion

On test day, mice were humanely euthanized 90 min following injection of either saline (placebo condition) or a drug (i.e., morphine/naloxone/opioid receptor agonist/CNO). This timing was selected to correspond with the peak expression of the immediate early gene product c-Fos, which was used as a measure of recent neuronal activity[59]. Mice were deeply anesthetized with sodium pentobarbital (80 mg/kg, i.p.) and underwent transcardial perfusion with 0.1LM phosphate-buffered saline (PBS, pH = 7.4) followed by 4% (w/v) paraformaldehyde in PBS (pH 7.4). Brains were extracted and post-fixed in the same fixative for 24h before being stored in 15% or 30% sucrose in PBS with 0.05% sodium azide at 4°C until cryosectioning.

### Immunohistochemistry and c-Fos analysis

All brains were sectioned on a cryostat (Cryostar NX50, ThermoFisher Scientific, Waltham, MA) at a thickness of 40 µm. Serial sections were collected and stored at −20L°C in a cryoprotectant solution consisting of 30% (w/v) sucrose, 1% (w/v) polyvinylpyrrolidone, and 30% (v/v) ethylene glycol in 0.1 M PBS until use. Free-floating sections were labelled for c-Fos using immunohistochemistry to quantify brain-wide patterns of neuronal activation. Sections were first washed several times in 0.1 M PBS (pH 7.4; 6× for 10 min each) and then incubated in a solution of 0.3% (v/v) H_2_O_2_/PBS for 30 min to minimize endogenous peroxidase activity. Sections were then rinsed several times in 0.1 M PBS before being placed in a blocking buffer containing 5% (v/v) Normal Goat Serum, 1% (w/v) Bovine Serum Albumin, and 0.3% (v/v) Triton X-100 dissolved in 0.1 M PBS for 2Lhr at room temperature. After blocking, sections were incubated with a primary anti-rabbit polyclonal c-Fos antibody (1:15,000, Synaptic Systems, #226 008, Gottingen, Germany) in the same blocking buffer for 48 h at 4L°C. Subsequently, sections were washed with 0.1 M PBS and incubated with anti-rabbit biotinylated secondary antibody (1:2000, Vector Laboratories, BA-1000, Newark, CA) for 2hr at room temperature. The antibody complex was then visualised using a DAB substrate Kit (SK-4100; Vector Laboratories) following the manufacturer’s instructions. The sections were mounted onto charged glass slides (Fisher SuperFrost Plus, Fisher Scientific, Whitby, ON) and left to air dry overnight. Slides were dehydrated through an ascending series of alcohols, cleared in xylene, and coverslipped with Entellan mounting medium.

Images were acquired at 10× using either a Cytation 5 (Biotek Cytation 5, Agilent, Santa Clara, CA) or Olympus VS200 slide scanner (Olympus VS200 slide scanner, Evident Corporation, Tokyo, Japan). Acquisition and analysis parameters were held constant across imaging systems, and cohorts were balanced across platforms. c-Fos expression was analyzed in 56 brain regions (Supplementary Table 1) where borders of regions were manually defined using the Franklin and Paxinos mouse brain atlas[60]. Fos-positive nuclei were quantified using

Image J software (National Institute of Health). Images were calibrated using pixels-to-distance ratio and were converted to 8-bit grey scale images. Fos-positive nuclei were detected and quantified using a consistent threshold across regions and images by experimenters who were blind to the condition. Three evenly spaced sections along the anterior-to-posterior axis were quantified bilaterally for most brain regions excluding the most anterior and posterior sections for each region to avoid sampling outside the region of interest. For a small subset of regions (e.g., dorsal raphe) where the anterior-posterior length of the region was less than 800 μm, counts were based on less than 3 sections. The quantification results were expressed as the average number of c-Fos-positive cells per standard unit of area (mm^2^).

For DREADD validation experiments, mice were perfused 90 min following CNO administration. Brains were cut into a 1:4 series. One series was dry mounted onto slides for histological reconstruction of the viral injection site. A second series underwent c-Fos immunofluorescent labelling. Sections were washed, incubated in the same c-Fos antibody (1:3000 in PBST for 72h at 4°C), and then visualised using a fluorophore-conjugated secondary antibody (donkey anti-rabbit Dylight488, ThermoFisher Scientific, Waltham, MA; 1:750 in PBST for 2h at room temperature). Sections were counterstained with DAPI, mounted with Aqua-Poly/Mount (Polysciences, Niles, Illinois, USA) antifade mounting medium, and imaged at 10× magnification using Olympus VS200 slide scanner. c-Fos-positive cells were quantified within predefined regions of interest corresponding to the ACC, BMA, and PVT. For each region, c-Fos expression was quantified across the anterior–posterior axis relative to the injection-site target coordinates by analysing 9–11 sections from the 1-in-4 series of 40 µm coronal sections, corresponding to approximately 1.3–1.6 mm of tissue centred on the target injection site. Regions of Interest (ROIs) were manually delineated in each section for each brain region and the number of c-Fos positive nuclei were then quantified using QuPath cell detection. Experimenters were blinded to condition during analysis.

### c-Fos Functional Connectivity

To assess coordinated activity across brain regions, whole-brain functional connectivity was quantified from inter-regional correlations in c-Fos density. For each experimental group a correlation matrix was generated using Pearson’s correlation coefficients (r) computed across animals for all included regions of interest. Correlation matrices were generated in GraphPad Prism 9 (GraphPad Software, Boston, MA, USA), with each matrix element representing the strength of the relationship between a pair of regions. Reliable estimation of inter-regional correlations required sufficient variability in c-Fos expression across animals. Therefore, datasets or regions with consistently low or zero values were excluded from the functional connectivity analysis. Home cage control animals were not included due to low overall c-Fos expression across the brain, resulting in a high proportion of zero or near-zero values. In addition, individual ROIs in which c-Fos density values were zero in more than two animals were excluded to avoid biasing correlation estimates. These regions were the lateral amygdala (LA), intercalated amygdala (IA), rhomboid thalamus (RH), ventral posterolateral thalamus (VPL), ventromedial hypothalamus (VMH), and posterior hypothalamus. The remaining ROIs were included in all subsequent analyses.

To enable comparison across conditions, network density was held constant by selecting the top 100 strongest correlations within each group. Comparisons of connection class distributions (e.g., cortex–cortex vs subcortex–subcortex) were therefore based on proportional differences rather than absolute edge counts. These connections were used to generate adjacency matrices in Microsoft Excel, representing the most robust functional interactions within each group. This approach ensured matched network density across conditions while allowing differences in connectivity structure to emerge without imposing a shared correlation threshold. These thresholded networks were used for visualization only and were not used for statistical comparisons. Functional connectivity networks were visualized using Gephi software (v0.10.1). Adjacency matrices were imported as undirected graphs, with nodes representing individual brain regions and edges representing functional connections. Networks were arranged using a circular layout with nodes ordered by anatomical identity. Node degree, defined as the number of connections per region within the thresholded network, was used as a measure of network centrality. For visualization purposes, brain regions were grouped into major anatomical categories (cortical regions, accumbens, hippocampus, amygdala, thalamus, hypothalamus, and brainstem), and nodes were colour-coded accordingly. Network graphs were exported and refined in Adobe Illustrator for figure preparation.

### Edgewise Functional Connectivity Comparisons

To identify condition-dependent differences in inter-regional coordination, functional connectivity was compared at the level of individual edges using the full, unthresholded correlation matrices. For each pair of regions, Pearson correlation coefficients were computed across animals within each condition and transformed using Fisher’s z-transformation prior to comparison. The empirical difference in correlation strength between the natural history and placebo groups was then calculated for each edge. Statistical significance was estimated using a permutation-based procedure implemented in R. Group labels were randomly permuted across animals without replacement, correlation matrices were recomputed, and between-group differences were recalculated for each permutation to generate a null distribution for each edge. This procedure was repeated 10,000 times. Observed differences were compared against the corresponding null distributions to obtain p-values, which were corrected for multiple comparisons using false discovery rate procedures to yield q-values. Significant edges were visualized on network graphs, with red edges indicating stronger connectivity and blue edges indicating weaker connectivity relative to the comparison group.

#### Regional connectivity strength

To assess the regional connectivity change among conditions, we calculated the normalized signed strength and its positive and negative components. Given a node (region) *i*, its strength *s* is defined as the sum of all correlation coefficients of node *i*. We calculated the positive 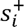 and negative 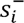 strengths and normalized them as

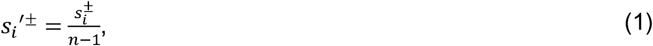

where *n* is the number of correlations. The normalized positive and negative strengths vary in the range of [0,1]. Next, we calculated the normalized signed strength such that positive contributions have more weight than negative ones as proposed in Rubinov & Sporns[61].

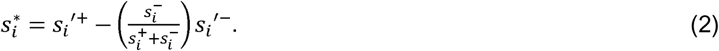

The 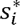 has a [-1,1] range. Then, we compared the 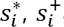 and 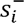 of each region among the conditions as described above. Group differences in positive strength, negative strength, and signed strength were assessed using the same permutation-based framework described above, with group labels permuted across animals and null distributions generated over 10,000 iterations. Resulting p-values were corrected for multiple comparisons using false discovery rate procedures across regions.

### Stereotaxic surgeries

All surgeries were performed under sterile conditions. Mice were anesthetized with isoflurane (5% induction, 2-4%% maintenance in oxygen) and secured in a stereotaxic frame. The scalp was sterilized, shaved, and the skull leveled using a digital stereotaxic instrument (Harvard Apparatus, Holliston, MA). A midline scalp incision was made to expose the skull, and small craniotomy holes (∼1 mm diameter) were drilled above target coordinates derived from a mouse brain atlas[60], taking care to avoid penetrating the dura mater. Microinjections of AAV viruses, dermorphin–saporin (derm-sap), or saporin control were performed using the following coordinates relative to bregma: anterior cingulate cortex (ACC), AP +1.20 mm, ML ±0.40 mm, DV −0.80 mm; basomedial amygdala (BMA), AP −1.55 mm, ML ±2.75 mm, DV −5.50 mm; paraventricular nucleus of the thalamus (PVT), AP −1.4 mm, ML +0.5 mm, DV −3.20 mm with a 10° angle. For chemogenetic manipulations, bilateral injections of AAV constructs (300 nl total at 50 nl/min) were performed using control virus (pAAV-CaMKIIα-mCherry, Addgene #114469; gift from Karl Deisseroth), inhibitory DREADD (pAAV-CaMKIIα-hM4D(Gi)-mCherry, Addgene #50477; gift from Bryan Roth), or excitatory DREADD (pAAV-CaMKIIα-hM3D(Gq)-mCherry, Addgene #50476; gift from Bryan Roth, Addgene, Watertown, MA, USA). For intersectional targeting of VGAT+/MOR+ neurons, VGAT-Cre mice received bilateral injections of a Cre- and Flp-dependent viral cocktail. This consisted of a Flp recombinase-expressing virus driven by the MOR promoter (pAAV-mMORp-FlpO, Stanford Neuroscience Gene Vector and Virus Core; gift from Karl Deisseroth Stanford University, Stanford, CA) and a CreON/FlpON excitatory DREADD construct (AAV-nEF-Con/Fon-hM3Dq-mCherry, Stanford Neuroscience Gene Vector and Virus Core; gift from Karl Deisseroth Stanford University). This strategy restricted DREADD expression to neurons co-expressing VGAT (Cre+) and MOR (Flp+). These viruses were co-infused bilaterally (1:1 ratio, 300nl/side at a rate of 60nl/min), and injectors were left in place for 5 min following infusion to allow diffusion. Viruses were allowed 3 weeks for expression prior to behavioral testing.

For neuronal ablation experiments, bilateral infusions of derm-sap or blank saporin (200 nl total; IT-12, KIT-12, Advanced Targeting Systems, Carlsbad, CA, USA) were delivered bilaterally at a rate of 50 nl/min using the ACC coordinates. Injectors were again left in place for 5 min prior to withdrawal. The scalp was closed with 6-0 coated Vicryl sutures (Ethicon, Somerville, NJ), and mice were returned to their home cages and administered postoperative analgesia (meloxicam, 20 mg/kg, every 24 h for three days). Behavioral assessment of placebo analgesia was performed 4 weeks after saporin injections.

### Verification of Viral Expression and Injection Sites

Histological validation of injection sites and subsequent viral expression was accomplished by 3D reconstruction of viral spread using the Histological E-data Registration in rodent Brain Spaces (HERBS) program. Brains were sectioned at 40 μm into a 1:4 series and dry mounted directly onto slides to preserve orientation and order, left to dry overnight, then incubated in DAPI (1:20000 in PBS) and cover slipped. Sections were imaged using VS200 Slidescanner and images were imported into QuPath, where the mCherry signal was thresholded using a fixed intensity criterion applied across all mice. For each injection site, eight sequential images were chosen centred around the injection target coordinates and were imported into the HERBS software[62]. Three-dimensional reconstructions of viral spread were generated from these serial coronal sections by aligning sections to Franklin and Paxinos reference atlas[60], and the extent of viral expression was manually outlined in each section. Outlines were then stacked across the anterior–posterior axis to generate volumetric reconstructions of injection spread. To ensure consistency across animals, all reconstructions were rendered using identical visualization parameters, including fixed scaling (width = 8), opacity (75%), and a uniform color mapping. Mice were excluded from analysis if viral expression was not detectable, was unilateral when bilateral targeting was required, missed the target ROI, or extended significantly beyond the anatomical boundaries of the target region. Fewer than 10% of animals were excluded based on these criteria.

### Western blotting

To assess μ-opioid receptor (MOR) protein levels, mice were euthanized by cervical dislocation, and brains were rapidly extracted, snap-frozen in liquid nitrogen, and stored at −80°C. Brains were sectioned into 1 mm coronal slices using a brain matrix, and ACC tissue was isolated by manual microdissection guided by stereotaxic coordinates corresponding to the injection site. Tissue was dissected on ice and homogenized in RIPA buffer (Millipore Sigma, 20-188) supplemented with protease and phosphatase inhibitors (Thermo Fisher Scientific, 78440), followed by centrifugation at 14,000 × g for 20 min at 4 °C. The supernatant was collected, and protein concentration was determined using a Bradford assay according to the manufacturer’s instructions (Bio-Rad Quick Start™ Bradford 1x Dye Reagent, 5000205). Equal amounts of protein (20 μg) were separated on 10% SDS-polyacrylamide gels (TGX™ FastCast™ Acrylamide Starter Kit, 10%, 1610172) using a Mini-PROTEAN® Tetra Vertical Electrophoresis Cell (Bio-Rad, 1658004) at a constant voltage of 250 V for 30 min, or until the dye front reached the bottom of the gel. Proteins were transferred onto polyvinylidene fluoride (PVDF) membranes using a wet transfer system (Mini Trans-Blot Module, 1703935). Membranes were blocked in 5% skim milk in Tris-buffered saline with 0.1% Tween-20 (TBS-T) for 1 h at room temperature. Membranes were incubated overnight at 4 °C with primary antibodies against MOR (Millipore Sigma, AB1580-I) and β-actin (Sigma-Aldrich, A5316-100UL) at a 1:5000 dilution in 3% bovine serum albumin (BSA) in TBS-T. Following incubation, membranes were washed three times (5 min each) with TBS-T and incubated with HRP-conjugated secondary antibodies (anti-rabbit: Cedarlane Labs, 711-035-152; anti-mouse: Cedarlane Labs, 515-035-062) for 1 h at room temperature. Signals were developed using chemiluminescent substrate (Thermo Scientific SuperSignal West Femto Maximum Sensitivity Substrate, 0034095) for 2 min and visualized with a chemiluminescence imaging system (iBright 1500; iBright 750). All samples were processed in parallel and imaged under non-saturating conditions. Band intensities were quantified using ImageJ within the linear range of detection and normalized to β-actin. MOR immunoreactivity was quantified as two predominant bands at approximately 50 and 70 kDa[63]. For each band, mean gray intensity was measured and local background immediately adjacent to the band was subtracted. Background-corrected MOR signals were normalized to β-actin from the same lane and expressed relative to a pooled reference sample containing protein from all experimental animals. Uncropped western blots, including the pooled reference lanes and all quantified samples, are shown in Supplementary Fig. 13.

### MERFISH/Allen data

Publicly available single-cell and spatial transcriptomic data from the Allen Brain Cell Atlas[64] were analyzed to characterize Oprm1 expression within the ACC. Neuronal subclasses were defined using the Allen reference taxonomy, and *Oprm1* expression was quantified across glutamatergic and GABAergic populations and across anterior–posterior axis. These analyses were used to define the distribution of *Oprm1* across molecularly defined neuronal populations and to guide interpretation of RNAscope findings.

### RNAscope In Situ Hybridization

RNAscope in situ hybridization was performed using the RNAscope Multiplex Fluorescent Reagent Kit v2 (Advanced Cell Diagnostics). Fresh-frozen brain sections (20 μm) were fixed, dehydrated, and treated with RNAscope Protease III prior to probe hybridization according to the manufacturer’s protocol. Target probes included Mm-*Oprm1* (ACD probe cat no. 493251), Mm-*Camk2a* (ACD probe cat no. 445231), and *Mm-Gad2* (ACD probe cat no. 439371). Following probe hybridization, signal amplification and fluorescent detection were performed according to kit instructions. Sections were counterstained with DAPI, coverslipped with ProLong Gold antifade mountant, and imaged at 10× using an Olympus VS200 slide scanner. Cells were classified as transcript-positive when ≥3 fluorescent puncta were detected within a DAPI-defined cell boundary. Co-expression was defined as the presence of target transcripts within the same cellular boundary. Quantification was performed across three sections per animal per bregma level and averaged per mouse prior to statistical analysis. Experimenters were blind to condition during analysis.

### Data Analysis and Statistics

Statistical analyses were performed using GraphPad Prism and R. Data are presented as mean

± SEM. Behavioral data were analyzed using one- or two-way ANOVA with repeated measures where appropriate, followed by Šídák or Tukey post hoc tests. All tests were two-sided unless otherwise specified. For network analyses, permutation-based testing (10,000 iterations) was used to assess significance, with false discovery rate correction applied for multiple comparisons. To calculate % anti-allodynia, the following formulae were used in a stepwise fashion:

**1. Trapezoid Method of Area Under the Curve (AUC) –** This formula calculates the area under the curve using a modified trapezoid method.

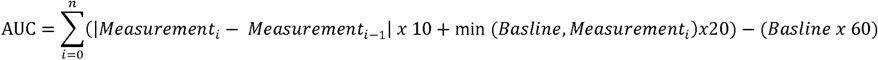
**2. Calculate % Allodynia**: This formula assesses the percentage increase in pain sensitivity by comparing the post-SNI baseline measurement to the pre-SNI baseline, reflected the extent of induced allodynia.

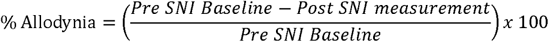
**3. Calculate % Anti-allodynia** – This formula was derived by comparing the adjusted AUC to the maximum AUC (trapezoid method without the adjustment), providing the percentage reduction in allodynia.

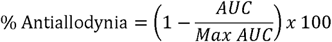

### Reproducibility

Experiments were performed in a minimum of three independent cohorts. Cohorts consisted of 6–10 mice (3–4 per experimental group) and were analyzed as independent replications, yielding consistent results across cohorts. Behavioral assessments and data analysis were conducted with experimenters blind to condition. c-Fos and immunohistochemical quantification were performed independently by at least two blinded experimenters.

## Data Availability

The data generated in this study are provided in the Supplementary Information/Source Data file. Source data are provided with this paper.

## Acknowledgments

We thank Hyun Been Park, Fiona Ramnaraign, Matthew Danesh, Areej Fatima, Harsukh Sidhu, Ahmed Aldarraji, Precia Christian, and Batul Presswala for assistance with early-stage c-Fos quantification. We also thank Claire Chan and Matthew Danesth and Sarasa Tohyama for early work on the placebo model

## Author Contributions

L.J.M. conceived and designed the study. C.C. initiated the project and contributed to data collection. S.K.R. and D.C.B. performed the majority of the experiments and collected the data. S.K.R. led the behavioural studies, including the MOR ablation, chemogenetic behavioural, and western blot experiments. D.C.B. performed viral injections and histological analyses and, together with L.J.M., led data analysis and figure preparation. C.A.O.C. performed the signed network analyses and contributed to their interpretation. S.A.K., J.B., F.K., S.R.G., and M.I.F. contributed to data collection and experimental procedures. P.W.F. and L.J.M. supervised the project and secured funding. S.K.R. and D.C.B. contributed to the preparation of the Methods, and L.J.M. wrote the manuscript with input from all authors. All authors discussed the results, revised the manuscript, and approved the final version.

