## Supplementary Figures for "Whole-brain analyses identify anterior cingulate μ-opioid signaling as a critical mediator of placebo analgesia in neuropathic pain"

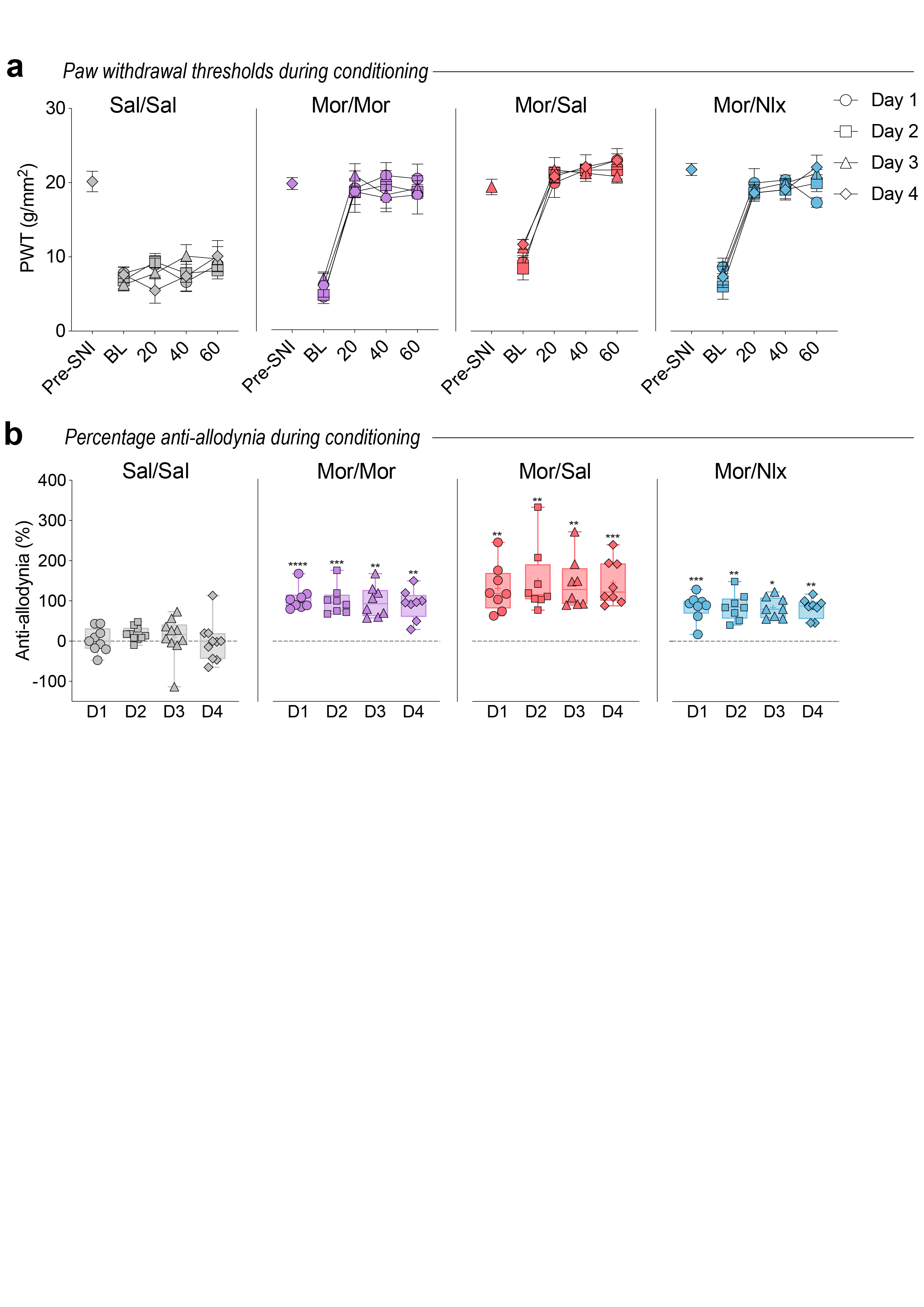

**Supplementary Figure 1. Comparable morphine-induced anti-allodynia across conditioning sessions prior to placebo testing.** **a** Paw withdrawal thresholds measured prior to nerve injury (Pre-SNI), immediately before drug administration on each conditioning day (BL), and at 20, 40, and 60 min following treatment. Symbols denote individual conditioning days. Statistical comparisons were performed on anti-allodynia values shown below. **b** Anti-allodynia calculated for each conditioning session. Each symbol represents an individual mouse, and symbol shape denotes conditioning day. Anti-allodynia differed between treatment groups but remained stable across conditioning sessions (two-way repeated-measures ANOVA, Group effect F(3,29) = 20.11, p < 0.0001; Time effect F(2.62,76.05) = 0.48, p = 0.67; Group × Time interaction F(7.87,76.05) = 0.32, p = 0.95). Post hoc comparisons revealed greater anti-allodynia in all morphine-treated groups relative to saline-treated controls on each conditioning day, whereas Mor/Sal, Mor/Mor, and Mor/Nlx groups did not differ from one another, indicating equivalent morphine conditioning prior to test-day manipulations. Data are presented as mean ± s.e.m. Individual data points are shown. **p < 0.01, ***p < 0.001, ****p < 0.0001 compared with Sal/Sal on the corresponding conditioning day.

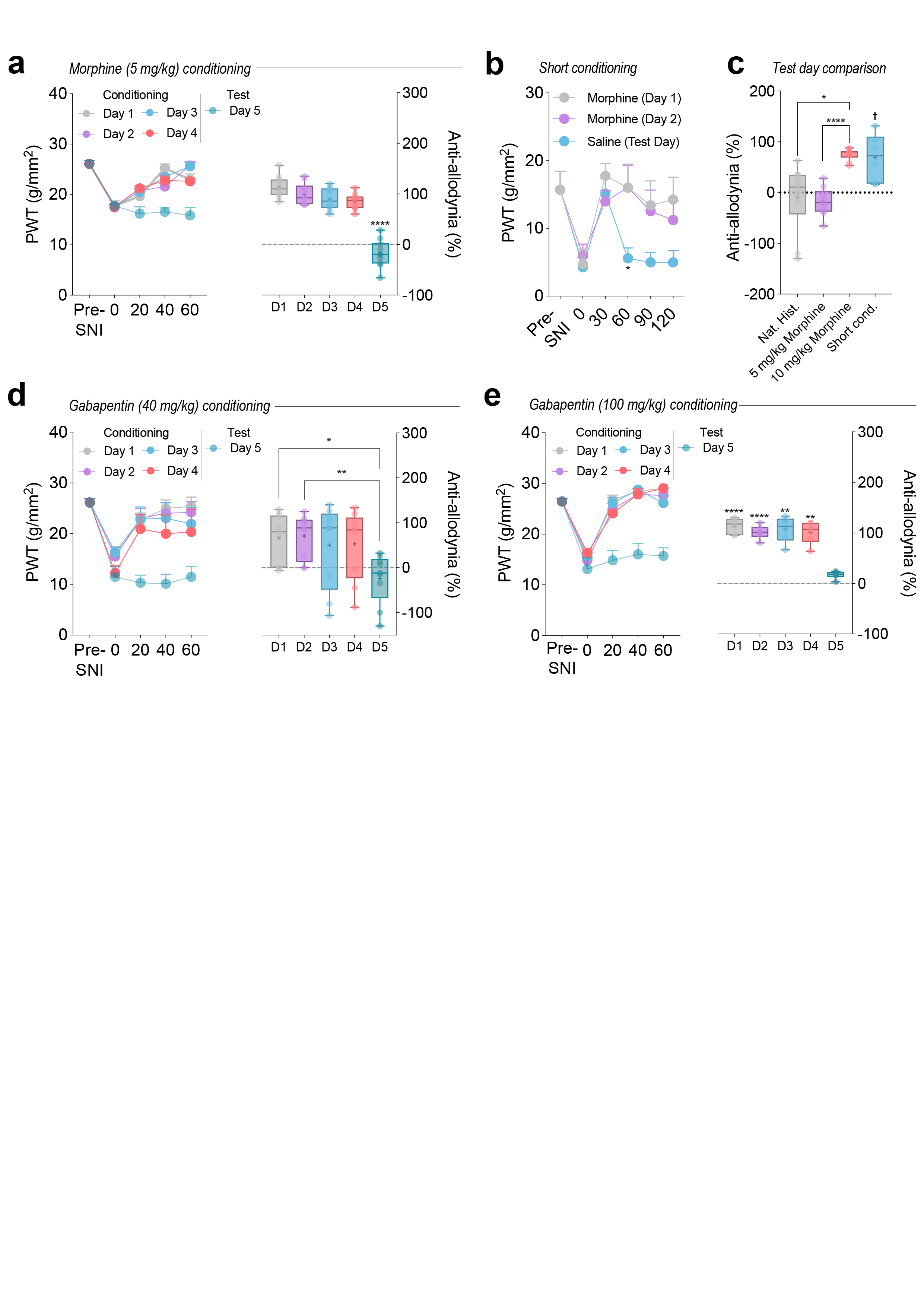

**Supplementary Figure 2. Placebo analgesia is not solely determined by conditioning-associated anti-allodynia.** **a** Left: Paw withdrawal thresholds measured prior to nerve injury (Pre-SNI), immediately before treatment (0 min), and 20, 40, and 60 min following treatment during each conditioning session and placebo test day. Statistical comparisons were performed on anti-allodynia values. Right: Anti-allodynia calculated for each conditioning session and placebo test day. Anti-allodynia differed across conditioning and test days (repeated-measures ANOVA, F(2.08,18.70) = 87.70, p < 0.0001). Post hoc Dunnett's comparisons demonstrated that anti-allodynia on the placebo test day (D5) was significantly reduced relative to D1 (adjusted p = 4.37 × 10⁻⁶), D2 (adjusted p = 1.39 × 10⁻⁶), D3 (adjusted p = 7.26 × 10⁻⁷), and D4 (adjusted p = 1.04 × 10⁻⁵). **b** Paw withdrawal thresholds measured before SNI surgery (Pre-SNI), immediately prior to injection (0 min), and 30, 60, 90, and 120 min following morphine administration during conditioning (Days 1–2; 10 mg/kg, i.p.) or saline administration on the placebo test day (Day 3). Two-way repeated-measures ANOVA revealed significant effects of time (F(2.78,13.92) = 7.23, p = 0.0042) and day (F(1.40,6.98) = 7.21, p = 0.0255), with no day × time interaction (F(2.44,12.22) = 1.70, p = 0.222). Post hoc Tukey's multiple comparisons tests identified lower withdrawal thresholds on the placebo test day relative to both conditioning days at 60 min following injection (Day 1 vs. Test Day, p = 0.0388; Day 2 vs. Test Day, p = 0.0164). **c** Comparison of placebo-induced anti-allodynia on the test day following natural history, 5 mg/kg morphine conditioning, 10 mg/kg morphine conditioning, or a short conditioning protocol. Anti-allodynia was calculated as the percentage reversal of SNI-induced mechanical hypersensitivity relative to baseline. Group differences were assessed using Brown–Forsythe and Welch ANOVA followed by Dunnett’s T3 multiple comparisons tests. Placebo responding differed across conditioning protocols, with placebo analgesia observed following 10 mg/kg morphine conditioning and the short conditioning protocol, whereas neither natural history controls nor 5 mg/kg morphine conditioning produced significant placebo analgesia. Placebo responding following the short conditioning protocol was comparable to that observed following 10 mg/kg morphine conditioning but significantly greater than that observed following 5 mg/kg morphine conditioning. Data are presented as box-and-whisker plots with individual animals shown. *p = 0.0132, ****p = 0.00000215, †p = 0.0217 compared with 5 mg/kg morphine. **d** Gabapentin conditioning produces transient anti-allodynia that is not maintained on the saline test day. Left panel shows paw withdrawal thresholds (PWTs) measured before nerve injury (Pre-SNI), immediately prior to drug administration (0 min), and 20, 40, and 60 min following gabapentin (40 mg/kg, i.p.) administration during conditioning (Days 1–4) or saline administration on the test day (Day 5). Right panel shows percentage anti-allodynia relative to pre-injection baseline for each day. Repeated-measures ANOVA revealed a significant effect of day (F(1.26,10.08) = 7.98, p = 0.014). Post hoc Dunnett's tests comparing the saline test day (D5) with each conditioning day demonstrated significantly lower anti-allodynia on D5 relative to D1 (p = 0.015) and D2 (p = 0.009), whereas D5 did not differ from D3 (p = 0.155) or D4 (p = 0.095). **e** Higher-dose gabapentin conditioning fails to produce placebo analgesia on the saline test day. Left panel shows paw withdrawal thresholds (PWTs) measured before nerve injury (Pre-SNI), immediately prior to drug administration (0 min), and 20, 40, and 60 min following gabapentin administration (100 mg/kg, i.p.) during conditioning (Days 1–4) or saline administration on the test day (Day 5). Right panel shows percentage anti-allodynia relative to pre-injection baseline for each day. Repeated-measures ANOVA revealed a significant effect of day (F(2.31,11.54) = 45.15, p = 2.34 × 10⁻⁶). Post hoc Dunnett's tests comparing the saline test day (D5) with each conditioning day demonstrated significantly lower anti-allodynia on D5 relative to D1 (p = 0.00021), D2 (p = 0.00046), D3 (p = 0.00154), and D4 (p = 0.00124). Time-course data are presented as mean ± s.e.m. Box-and-whisker plots show the median, interquartile range, and minimum-to-maximum values, with individual animals overlaid. Anti-allodynia was calculated as the percentage reversal of SNI-induced mechanical hypersensitivity relative to pre-injury baseline. Exact p values are reported in the text.

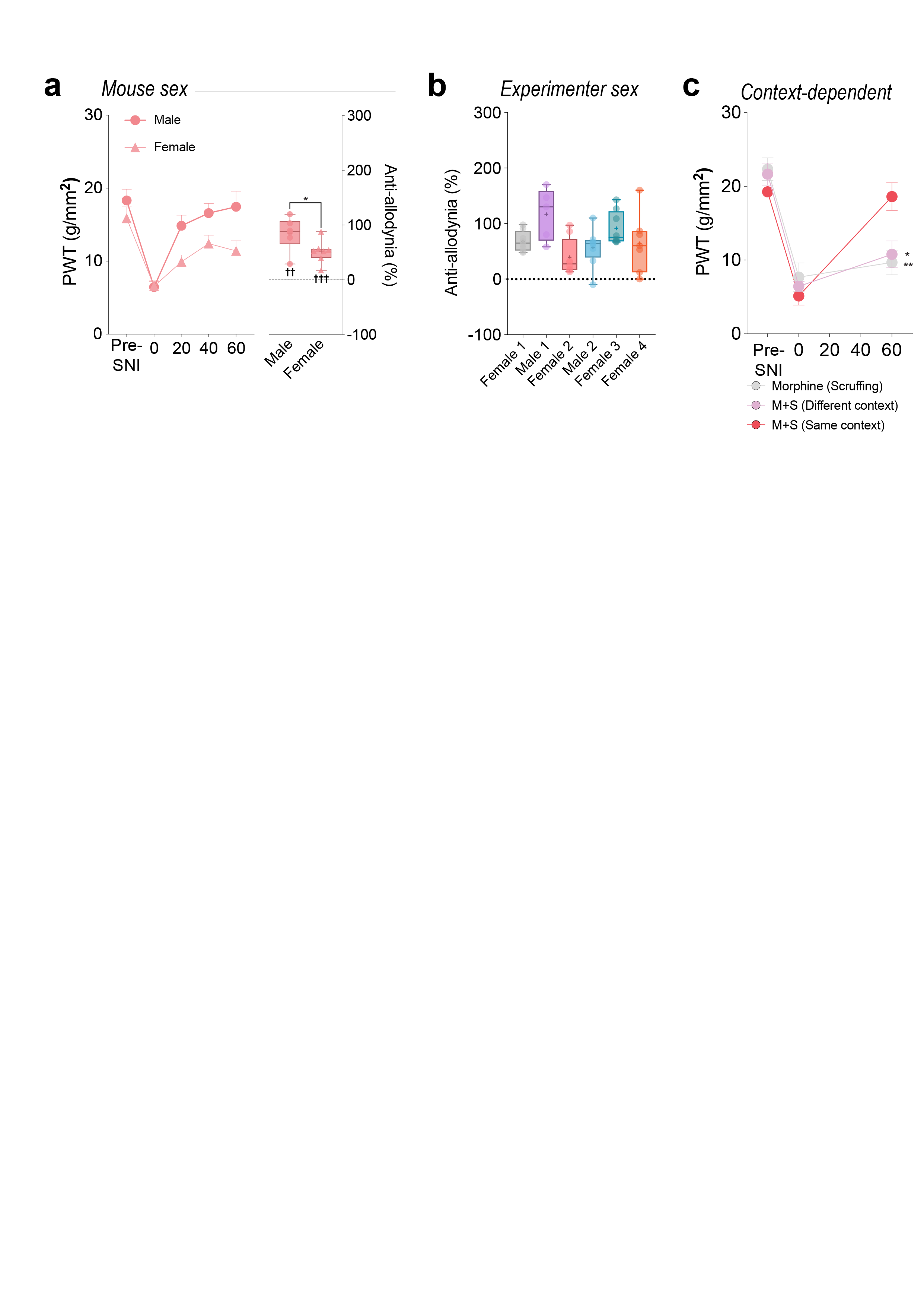

**Supplementary Figure 3. Placebo analgesia is reproducible across mouse sex, multiple experimenters and depends on contextual re-exposure.** **a** Placebo analgesia is observed in both male and female mice. Left: Paw withdrawal thresholds (PWTs) measured before nerve injury (Pre-SNI), immediately prior to saline administration (0 min), and 20, 40, and 60 min following placebo testing. Right: Percentage anti-allodynia relative to pre-injury baseline. Both male and female mice exhibited significant anti-allodynia relative to zero during placebo testing (one-sample t-tests, males: t₅ = 6.67, p = 0.0011; females: t₆ = 6.49, p = 0.0006). Anti-allodynia was greater in male mice than in female mice (unpaired two-tailed t-test, t₁₁ = 2.26, p = 0.045). **b** Placebo-associated anti-allodynia across independent cohorts tested by multiple blinded male and female experimenters. Anti-allodynia was calculated as the percentage reversal of SNI-induced mechanical hypersensitivity relative to baseline. Significant placebo-associated anti-allodynia was observed in all cohorts (one-sample t-tests: F1, t₇ = 10.46, p = 1.59 × 10⁻⁵; M1, t₄ = 5.61, p = 0.00495; F2, t₇ = 3.42, p = 0.0112; M2, t₇ = 4.68, p = 0.00227; F3, t₇ = 8.57, p = 5.85 × 10⁻⁵; F4, t₆ = 3.22, p = 0.0182). **c** Mice previously conditioned with morphine were tested under one of three conditions: handling alone (Morphine (Scruffing)), exposure to a different context (M+S Different context), or re-exposure to the original morphine-paired context (M+S Same context). Two-way repeated-measures ANOVA revealed a significant time × condition interaction (F(3.31,33.06) = 5.42, p = 0.0030), a significant effect of time (F(1.65,33.06) = 65.84, p < 0.0001), and no main effect of condition (F(2,20) = 0.98, p = 0.393). Post hoc Tukey's tests showed that withdrawal thresholds remained significantly lower than pre-injury baseline at 60 min in mice exposed to handling alone (BL vs. 60 min, p = 0.0053) or a different context (BL vs. 60 min, p = 0.020), whereas withdrawal thresholds in mice re-exposed to the original conditioning context did not differ from baseline (BL vs. 60 min, p = 0.954). In the same-context group, withdrawal thresholds increased between 0 and 60 min (p = 0.00034). Data are presented as mean ± s.e.m. (line graphs) or box-and-whisker plots showing the median, interquartile range, and minimum-to-maximum values, with individual animals overlaid. In panel a, † indicates significant anti-allodynia relative to zero (one-sample t-tests). In panel c, *p < 0.05 and **p < 0.01 indicate significant differences between the 60 min time point and pre-injury baseline within each condition (Tukey's multiple comparisons tests).

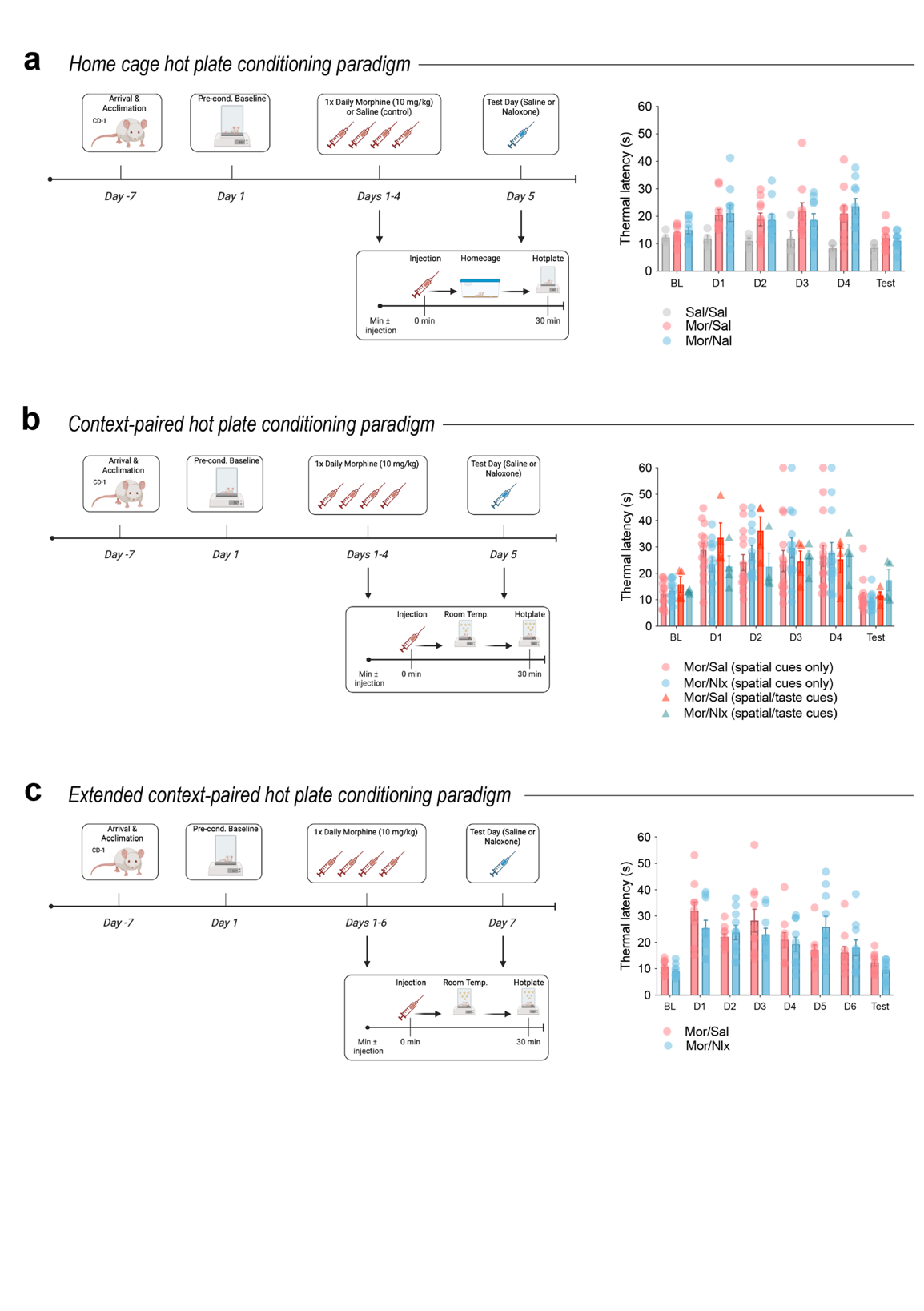

**Supplementary Figure 4. Repeated morphine conditioning does not produce placebo-like thermal analgesia in non-injured mice.** **a** Home-cage hot plate conditioning paradigm. Mice received morphine (10 mg/kg) or saline for four consecutive days and were tested on the hot plate 30 min later. Morphine-conditioned mice exhibited increased thermal withdrawal latencies during the conditioning phase, confirming successful conditioning. A two-way repeated-measures ANOVA revealed significant effects of time (F(3.41,71.67) = 6.16, p = 0.0005) and condition (F(2,21) = 3.94, p = 0.035), but no time × condition interaction (F(6.83,71.67) = 1.54, p = 0.168). Thermal withdrawal latencies did not differ between groups on the test day and were not altered by naloxone administration, indicating an absence of placebo analgesia. **b** Context-dependent hot plate conditioning using spatial cues alone or combined spatial and taste cues. Mice received morphine (10 mg/kg) paired with the conditioning context for four consecutive days and were tested on the hot plate 30 min later. Thermal withdrawal latencies increased during conditioning but returned to baseline levels on the test day. A two-way repeated-measures ANOVA revealed a significant effect of time (F(2.67,82.70) = 17.29, p < 0.0001), but no effect of condition (F(3,31) = 0.26, p = 0.855) or time × condition interaction (F(8.00,82.70) = 1.13, p = 0.353), indicating that neither conditioning procedure nor naloxone treatment produced placebo analgesia. **c** Extended hot plate conditioning paradigm. Mice received six days of morphine-context pairings before placebo testing. Thermal withdrawal latencies increased during the conditioning phase and returned to baseline levels on the test day. A two-way repeated-measures ANOVA revealed a significant effect of time (F(3.76,67.61) = 22.48, p < 0.0001) and a time × condition interaction (F(3.76,67.61) = 2.92, p = 0.030), but no main effect of condition (F(1,18) = 0.07, p = 0.796). Saline- and naloxone-treated mice did not differ on the test day (p = 0.114), indicating that extending the conditioning protocol did not produce detectable placebo analgesia. Data are presented as mean ± SEM. Symbols represent individual mice.

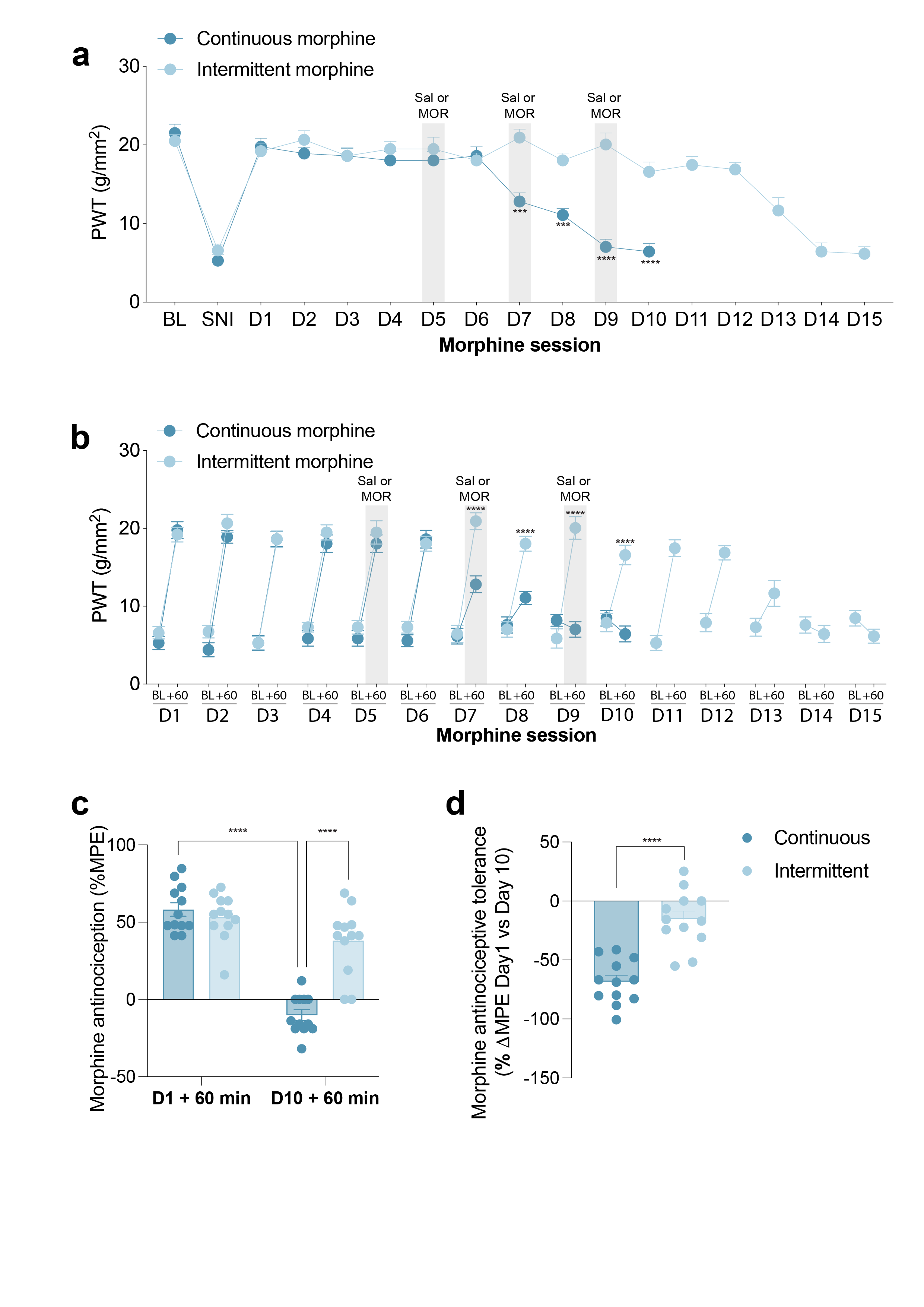

**Supplementary Figure 5. Intermittent saline administration delays the development of morphine tolerance in mice with spared nerve injury.** **a** Intermittent saline administration delays the development of morphine tolerance in mice with spared nerve injury. a Mechanical paw withdrawal thresholds (PWTs) measured before surgery (BL), following spared nerve injury (SNI), and during repeated morphine administration. Mice received either continuous daily morphine or intermittent morphine, in which saline was substituted for morphine on days 5, 7, and 9 (grey shaded regions). A two-way repeated-measures ANOVA (BL–D10) revealed significant effects of time (F(7.58,166.79) = 40.38, p < 0.0001), treatment group (F(1,22) = 23.86, p = 0.00007), and a time × group interaction (F(7.58,166.79) = 12.22, p < 0.0001). Šídák post hoc comparisons identified significant differences between groups on D7 (p = 0.00029), D8 (p = 0.00019), D9 (p = 0.000005), and D10 (p = 0.00003). Data from D11–D15 are shown for the intermittent morphine group only. b Mechanical withdrawal thresholds measured immediately before (BL) and 60 min after morphine administration (+60 min) across repeated treatment sessions. Pre-morphine thresholds did not differ between groups (time × group interaction: F(6.00,131.91) = 1.35, p = 0.2407; group: F(1,22) = 0.64, p = 0.4331). Post-morphine thresholds exhibited significant effects of time (F(5.38,118.36) = 17.60, p < 0.0001), treatment group (F(1,22) = 24.95, p < 0.0001), and a time × group interaction (F(5.38,118.36) = 13.42, p < 0.0001). c Morphine antinociception expressed as maximal possible effect (MPE) following the first (D1) and last (D10) morphine administration. A two-way repeated-measures ANOVA revealed significant effects of time (F(1,22) = 90.68, p = 0.000000003), treatment condition (F(1,22) = 18.44, p = 0.000294), and a time × condition interaction (F(1,22) = 36.42, p = 0.0000045). d Change in morphine antinociception between D1 and D10 expressed as ΔMPE. Groups differed significantly (Welch's two-tailed t-test, t = 6.04, df = 20.85, p < 0.0001). Data are presented as mean ± SEM.

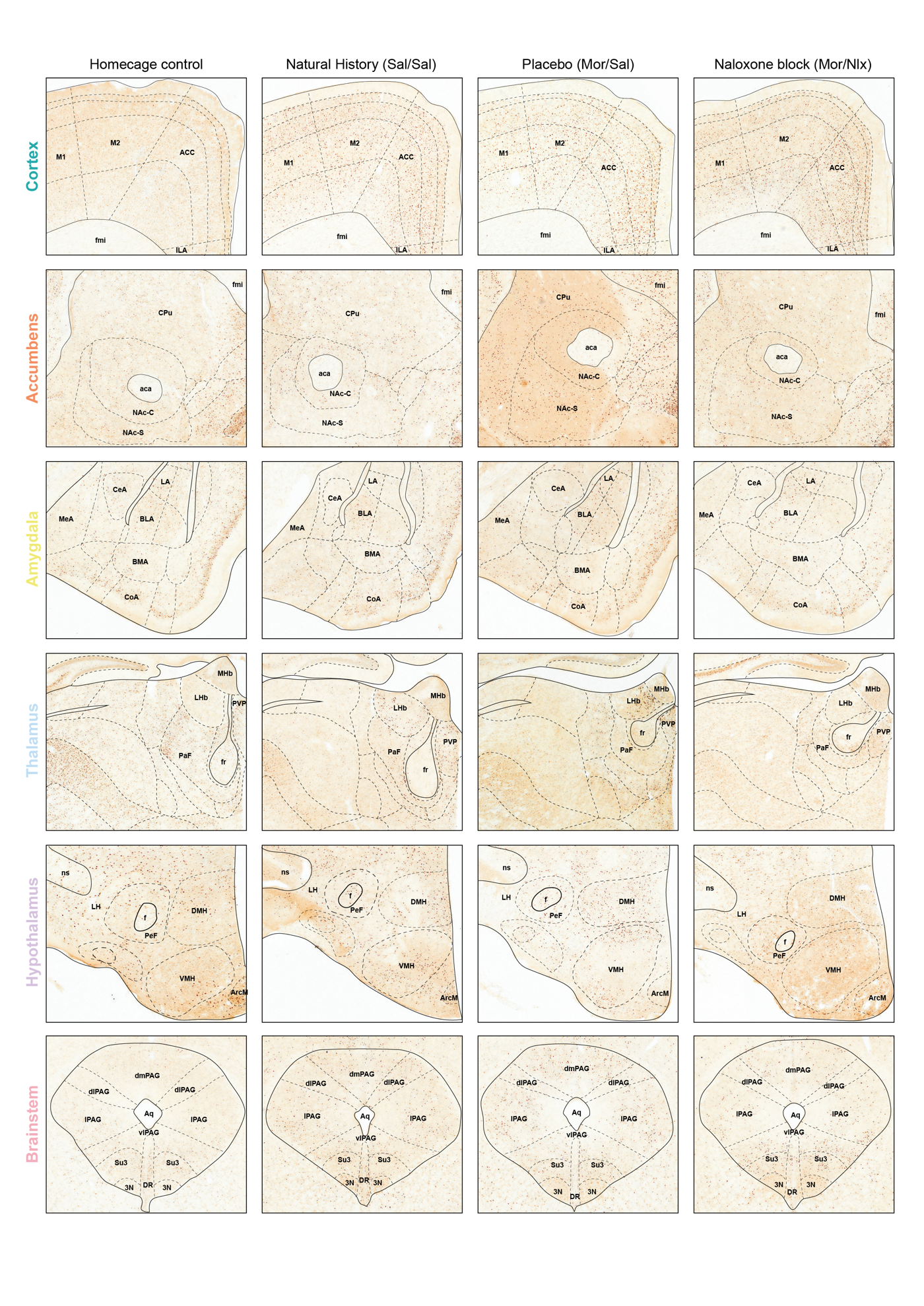

**Supplementary Figure 6. Representative c-Fos immunohistochemistry images from the whole-brain analysis.** Representative images from selected cortical, accumbal, amygdalar, thalamic, hypothalamic, and brainstem regions obtained from home cage, natural history (Sal/Sal), placebo (Mor/Sal), and naloxone-treated (Mor/Nlx) animals. Images are shown to illustrate representative staining patterns used for whole-brain c-Fos quantification. Corresponding quantitative analyses are presented in Figure 2.

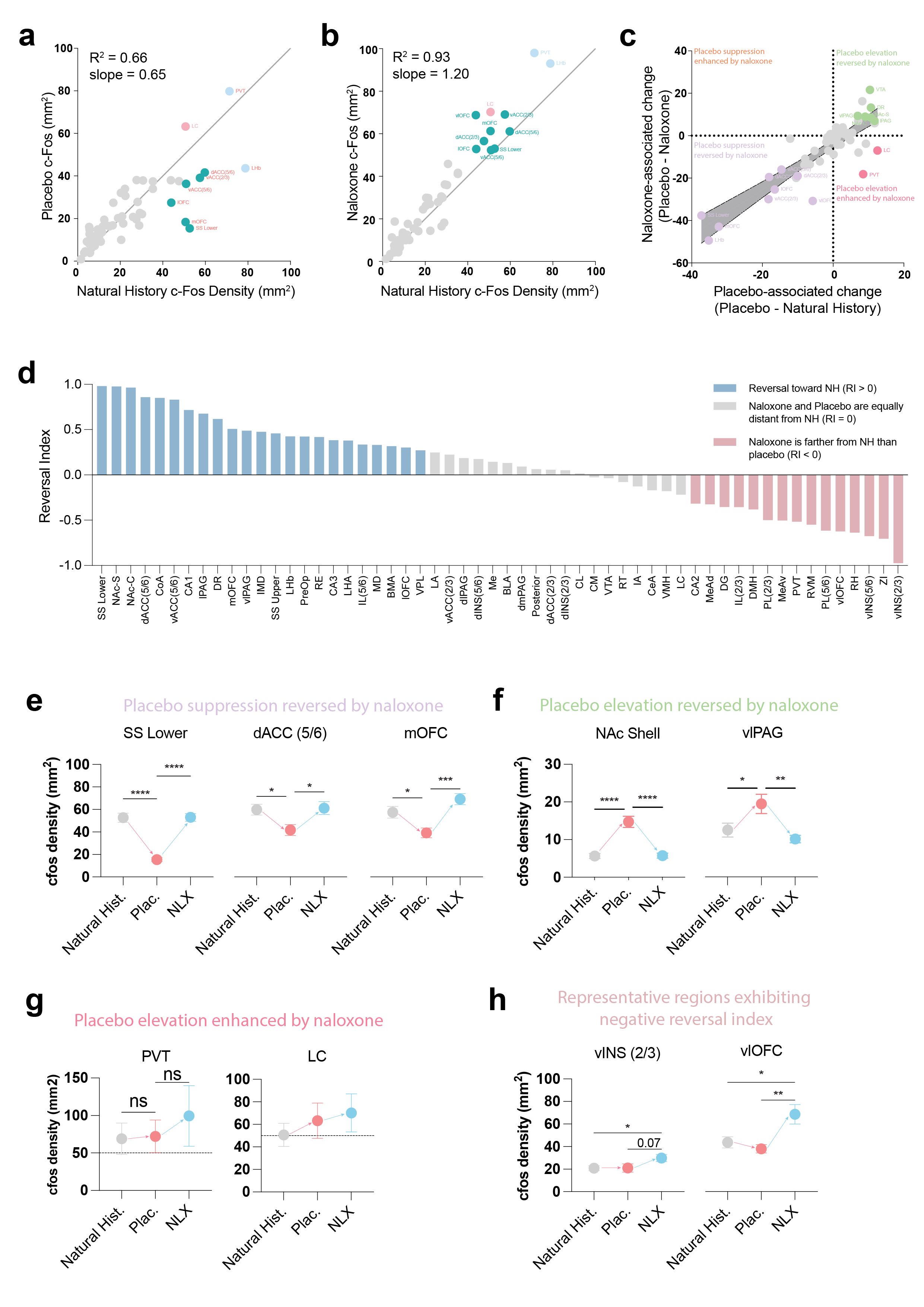

**Supplementary Figure 7. Naloxone selectively perturbs subsets of placebo-associated brain-wide c-Fos activity.** **a,b** Scatterplots showing regional c-Fos densities across quantified ROIs in placebo versus natural history mice (a) and naloxone-treated placebo versus natural history mice (b). Regional activity patterns remained strongly correlated across conditions. Highlighted ROIs indicate regions exhibiting prominent placebo-associated or naloxone-sensitive effects. Gray line indicates unity. **c** Relationship between placebo-associated regional changes and naloxone-induced shifts. ROIs in the upper-left and lower-left quadrants indicate naloxone-induced reversal toward natural history levels, whereas ROIs in the upper-right and lower-right quadrants indicate placebo-associated changes maintained following naloxone administration. Gray shading denotes the error bands surrounding the unity relationship. **d** Ranked naloxone reversal index across quantified ROIs. Positive values indicate reversal toward natural history levels, values near zero indicate persistence of placebo-associated activity, and negative values indicate divergence from the natural history state following naloxone administration. **e** Representative cortical regions exhibiting placebo-associated suppression reversed by naloxone, including lower-limb somatosensory cortex (SSLower; F(2,33) = 43.89, p < 0.0001), dorsal anterior cingulate cortex layer 5/6 [dACC (5/6); F(2,33) = 4.46, p = 0.0194], and medial orbitofrontal cortex (mOFC; F(2,33) = 9.75, p = 0.0005). **f** Representative regions exhibiting placebo-associated elevations reversed by naloxone, including nucleus accumbens shell (NAcShell; F(2,33) = 26.33, p < 0.0001) and ventrolateral PAG (vlPAG; F(2,33) = 6.49, p = 0.0042). **g** Representative regions exhibiting persistent placebo-associated elevations following naloxone administration, including paraventricular thalamus (PVT) and locus coeruleus (LC). **h** Representative regions exhibiting negative reversal indices, including ventral insular cortex layer 2/3 [vINS (2/3); F(2,55) = 2.67, p = 0.0785] and ventrolateral orbitofrontal cortex (vlOFC; F(2,33) = 7.04, p = 0.0028). Statistical comparisons in panels e–h were performed using one-way ANOVA with Šídák’s multiple-comparisons test.

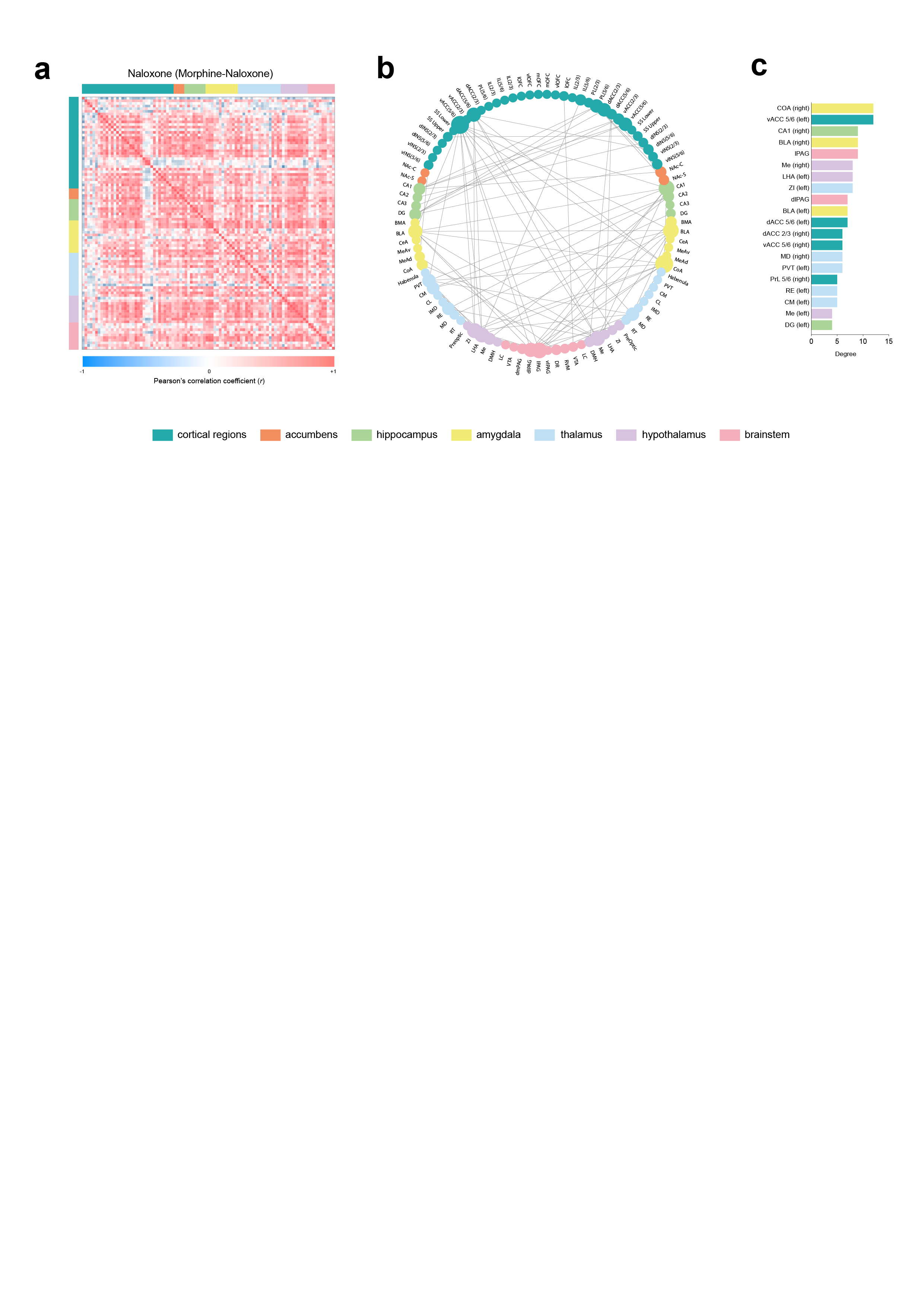

**Supplementary Figure 8. Whole-brain functional connectivity in the naloxone group.** **a** Group-level functional connectivity matrix for the naloxone group, generated from Pearson correlations of regional c-Fos densities across all quantified brain regions. Warmer colours indicate stronger positive correlations and cooler colours indicate weaker or negative correlations. Regions are ordered and colour-coded by major anatomical category, as in Figure 3. **b** Circular network representation of the top 100 strongest functional connections in the naloxone group. Nodes represent individual brain regions, grouped by anatomical category, and edges represent undirected functional connections. **c** Node degree distribution for the naloxone network, defined as the number of connections per region within the top-100 connection network. Bars are colour-coded according to anatomical group.

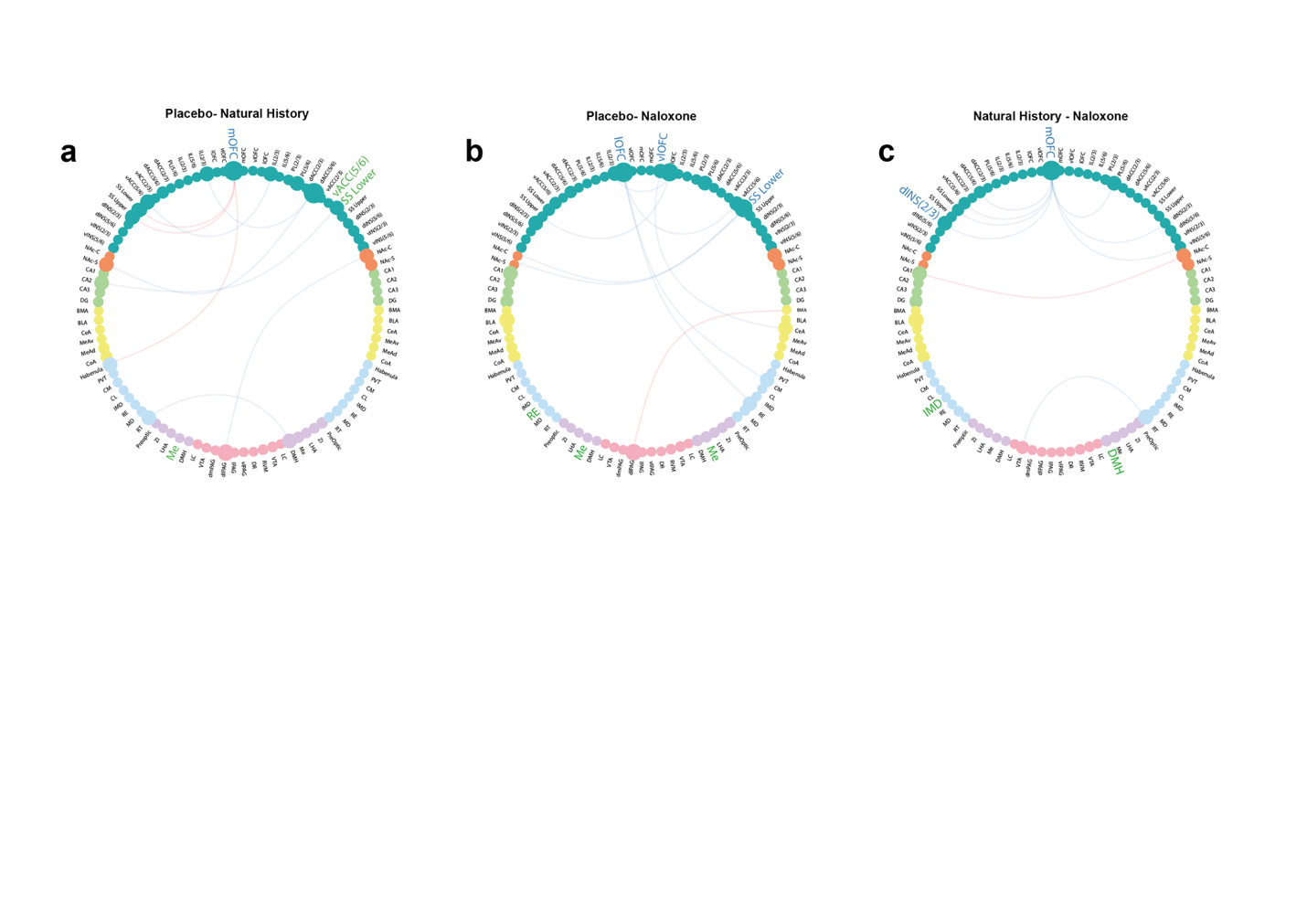

**Supplementary Figure 9.** Placebo analgesia and opioid receptor blockade are associated with distinct patterns of network reorganization across pain-related brain regions. Network maps illustrate significant differences in regional connectivity strength and pairwise functional correlations between Placebo and Natural History (**a**), Placebo and Naloxone (**b**), and Natural History and Naloxone (**c**). Relative to Natural History, placebo analgesia was associated with altered connectivity strength represented by the medial orbitofrontal cortex (mOFC), ventral anterior cingulate cortex [vACC (5/6)], and lower-limb somatosensory cortex (SS Lower), accompanied by widespread changes in functional connectivity (**a**). Comparison of Placebo and Naloxone networks identified differences involving orbitofrontal, somatosensory, reticular thalamic, hypothalamic, and amygdalar regions, represented by mOFC, ventrolateral orbitofrontal cortex (vlOFC), SS Lower, reticular thalamic nucleus (RE), medial hypothalamus (Me), and basomedial amygdala (BMA) (**b**). Direct comparison of Natural History and Naloxone networks revealed altered connectivity strength within orbitofrontal, insular, thalamic, and hypothalamic regions, represented by mOFC, ventral insular cortex [vINS (2/3)], intermediodorsal thalamus (IMD), and dorsomedial hypothalamus (DMH), together with corresponding alterations in pairwise functional connectivity (**c**). Nodes represent individual brain regions organized by anatomical grouping. Labeled regions exhibited significant differences in normalized connectivity strength metrics following permutation testing with false discovery rate correction. Blue labels indicate regions with greater connectivity strength in the first condition listed for each comparison, whereas green labels indicate regions with greater connectivity strength in the second condition. Connectivity strength differences could arise from alterations in positive connectivity strength (Spos), negative connectivity strength (Sneg), or signed strength. Edges represent pairwise correlations that differed significantly between conditions. Red edges indicate correlations that were significantly stronger in the first condition listed for each comparison, whereas blue edges indicate correlations that were significantly weaker in the first condition. Only statistically significant differences are shown.

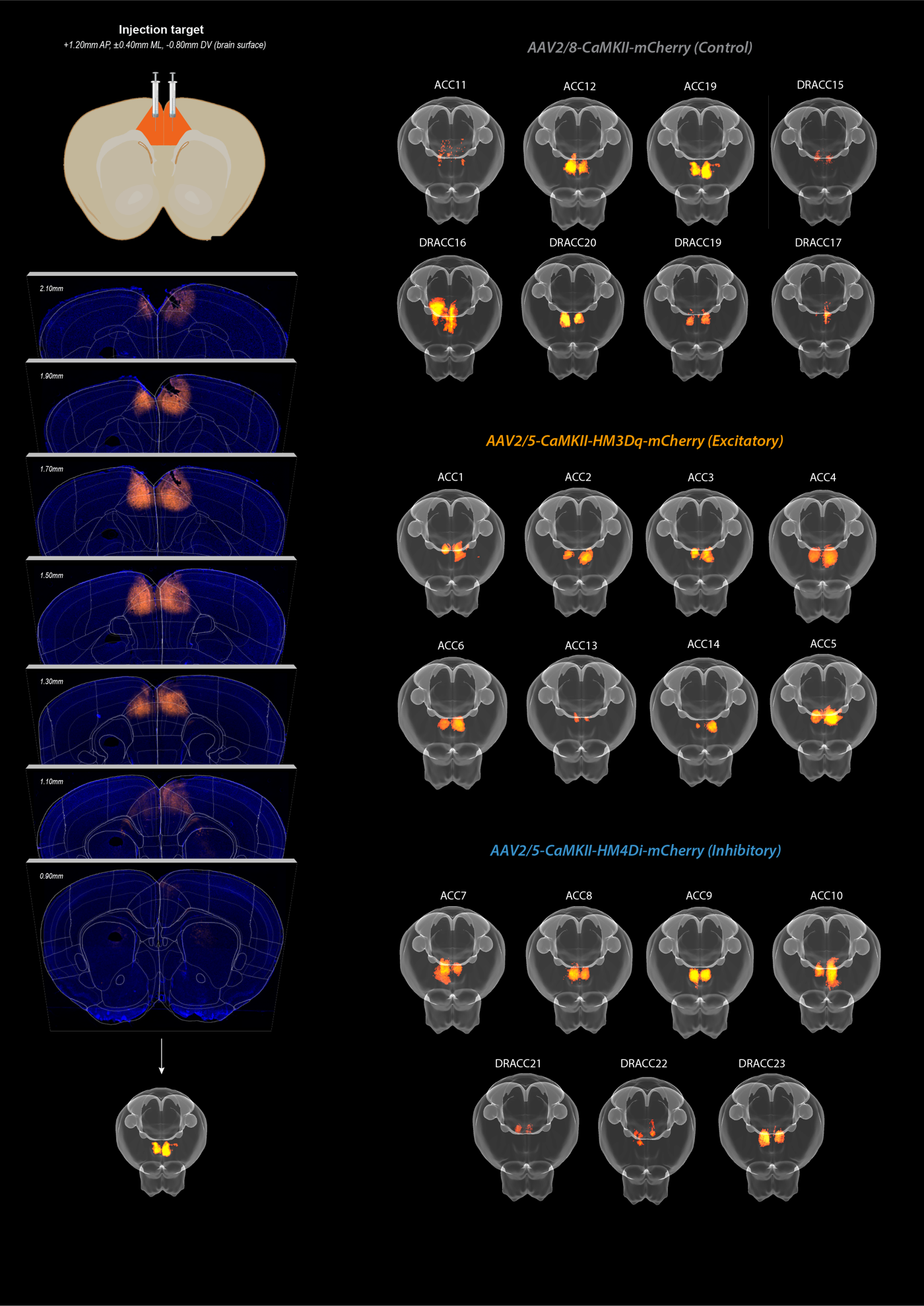

**Supplementary Figure 10. Verification of viral expression and injection targeting in the anterior cingulate cortex (ACC).** (Top left) Schematic showing the intended injection coordinates targeting the ACC at +1.20mm AP, ±0.40mm ML, and –0.8mm DV relative to brain surface. (Middle left) Histological verification of mCherry expression across coronal sections from a representative animal, shown at multiple anteroposterior levels spanning the ACC. (Bottom left) 3D reconstruction of mCherry expression from that animal using HERBS. (Right panels) Individual 3D reconstructions of mCherry expression for all included animals in each group are shown: AAV2/8-CaMKIIα-mCherry (Control, top), AAV2/5-CaMKIIα-hM3Dq-mCherry (Excitatory DREADD, middle), and AAV2/5-CaMKIIα-hM4Di-mCherry (Inhibitory DREADD, bottom). Each brain represents a distinct animal. Reconstructed volumes were aligned using HERBS software.

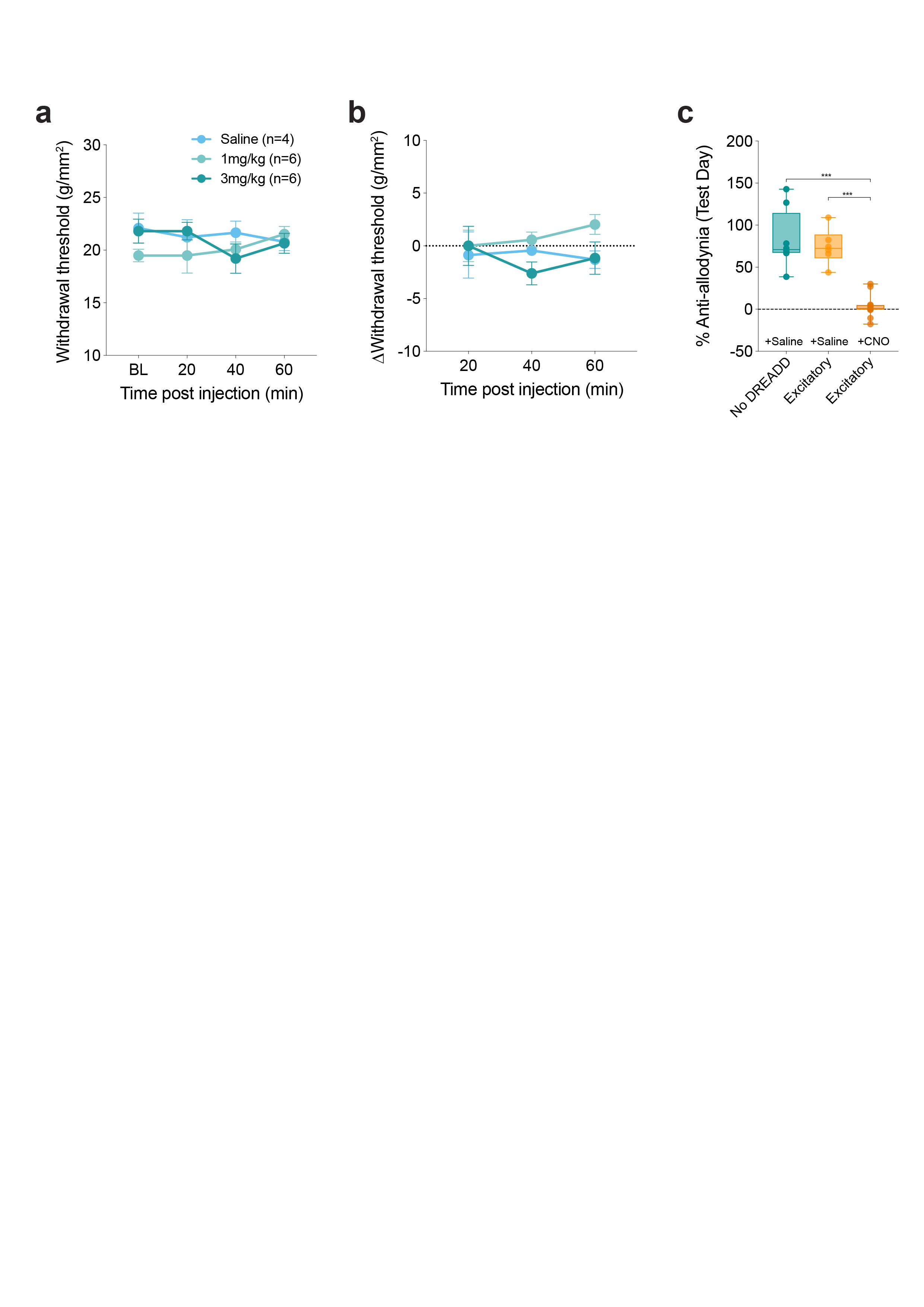

**Supplementary Figure 11. CNO administration does not alter baseline mechanical sensitivity but abolishes placebo analgesia following activation of excitatory ACC DREADDs.** **a** Mechanical withdrawal thresholds measured using the von Frey SUDO method (converted to g/mm²) pre-SNI before (BL; 0 min) and 20, 40, and 60 min following intraperitoneal injection of saline, CNO (1 mg/kg), or CNO (3 mg/kg) (time × treatment interaction (F(4.30,27.94) = 1.21, p = 0.330). These experiments were performed three weeks after AAV injection to match the timeline used for DREADD experiments. **b** Change in withdrawal threshold relative to baseline (Δ withdrawal threshold; g/mm²) for the same animals shown in (**a**), demonstrating no effect of systemic CNO administration on mechanical sensitivity in uninjured mice (time × treatment interaction (F(3.66,23.76) = 1.19, p = 0.340). **c** Saline-treated mice expressing hM3Dq DREADDs exhibited placebo responses comparable to saline-treated mice lacking DREADDs (p = 0.7820). CNO administration in hM3Dq-expressing mice abolished placebo-associated anti-allodynia, reducing responses relative to both saline-treated hM3Dq mice (p < 0.0001) and saline-treated mice lacking DREADDs (p < 0.0001; treatment effect: F(2,23) = 34.07, p < 0.0001). Groups: No DREADD + Saline (n = 8), Excitatory DREADD + Saline (n = 6), Excitatory DREADD + CNO (n = 12).. Data are shown as individual mice with mean ± SEM.

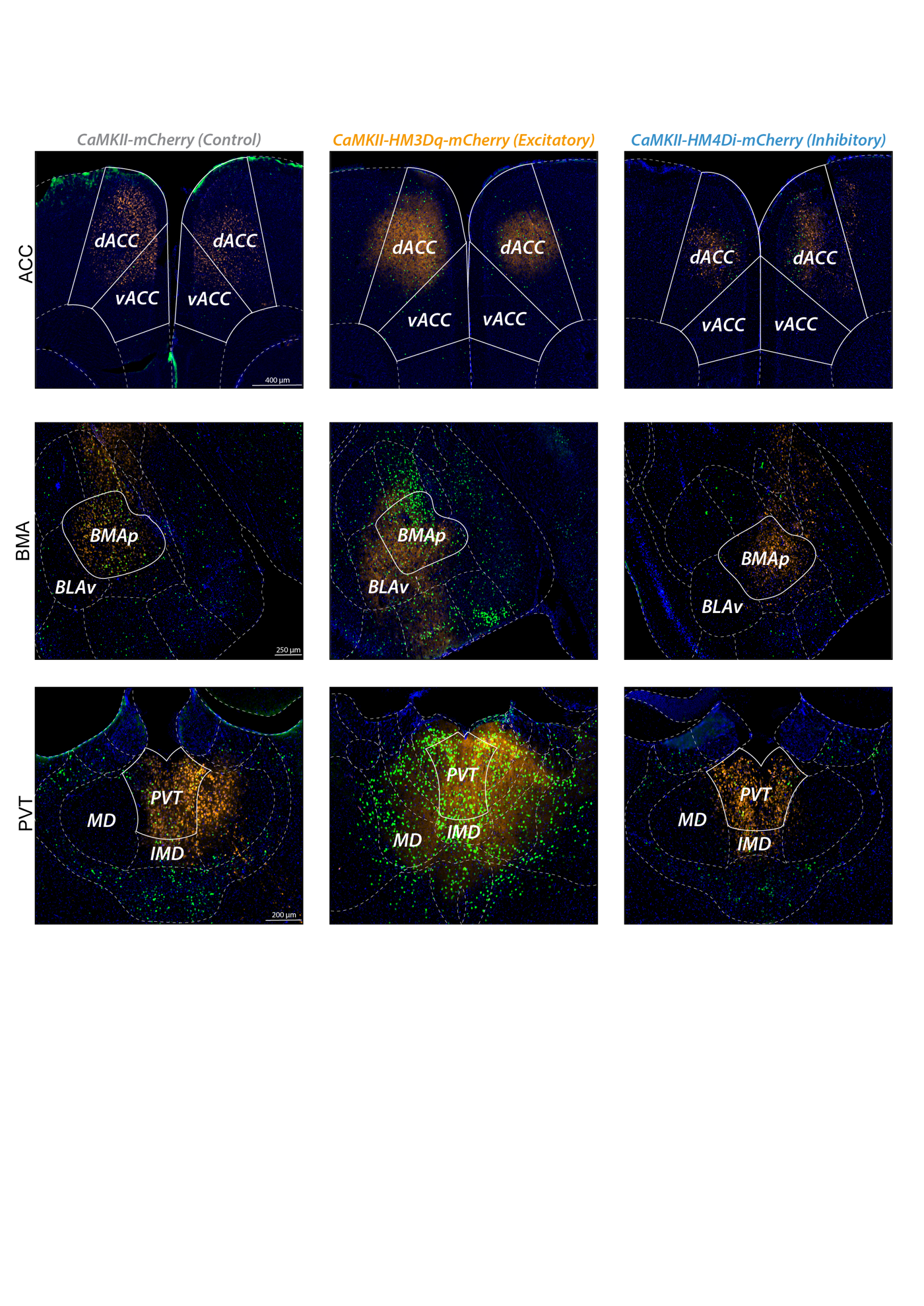

**Supplementary Figure 12. Representative c-Fos immunohistochemistry images from regions selected for chemogenetic manipulation experiments.** Representative images showing c-Fos expression within the anterior cingulate cortex (ACC; **top row**), basomedial amygdala (BMA; **middle row**), and paraventricular thalamus (PVT; **bottom row**) from Natural History, Placebo, and Naloxone mice. Images are shown to illustrate representative patterns of neuronal activity within these regions across control and DREADD conditions.

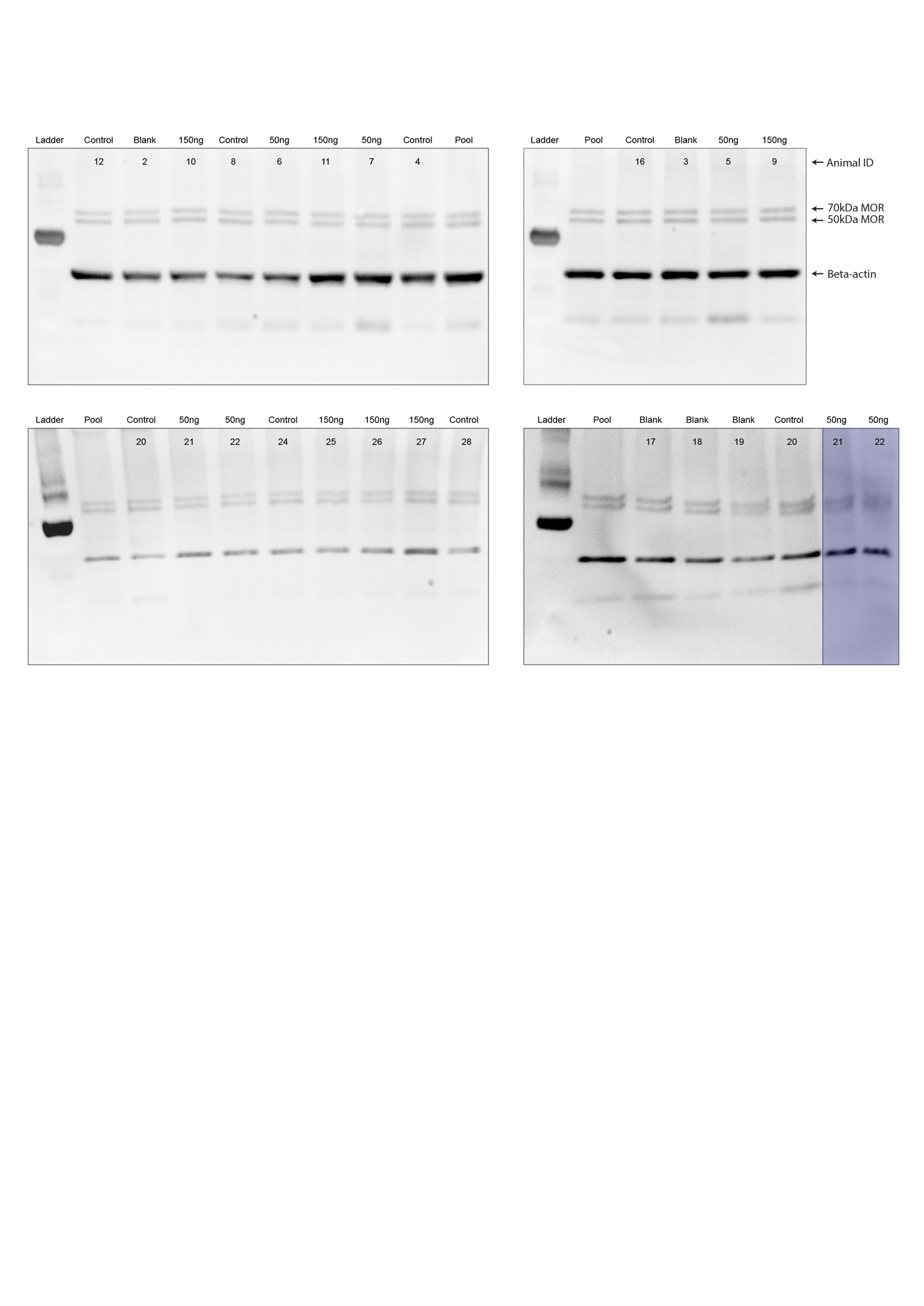

**Supplementary Figure 13. Full western blots confirming selective ablation of MOR+ neurons in the ACC following Dermorphin-Saporin injection.** Uncropped western blots of μ-opioid receptor (MOR) protein expression in anterior cingulate cortex (ACC) tissue collected one month following bilateral ACC injections of Dermorphin-Saporin (Derm-SAP). Mice received no injection (Control), blank saporin (Blank-SAP), 50 ng Derm-SAP, or 150 ng Derm-SAP. Four independent gels are shown. A pooled ACC sample (“Pool”) was included on each gel to allow normalization across membranes. Blots were probed using the same MOR and β-actin antibodies described in the main manuscript (Figure 6 and Methods). Two MOR-immunoreactive bands were detected at ~70 kDa and ~50 kDa, consistent with previously reported full-length and processed MOR species. β-actin served as a loading control. Densitometric quantification of the 70 kDa and 50 kDa MOR bands, normalized to β-actin and standardized across gels using the pooled control sample, is presented in Figure 6 of the main manuscript. All raw blots used for quantification are shown here. Two lanes (bottom right panel) exhibited technical artifacts and were repeated in a subsequent gel (bottom left panel); the repeated samples were used for quantification.

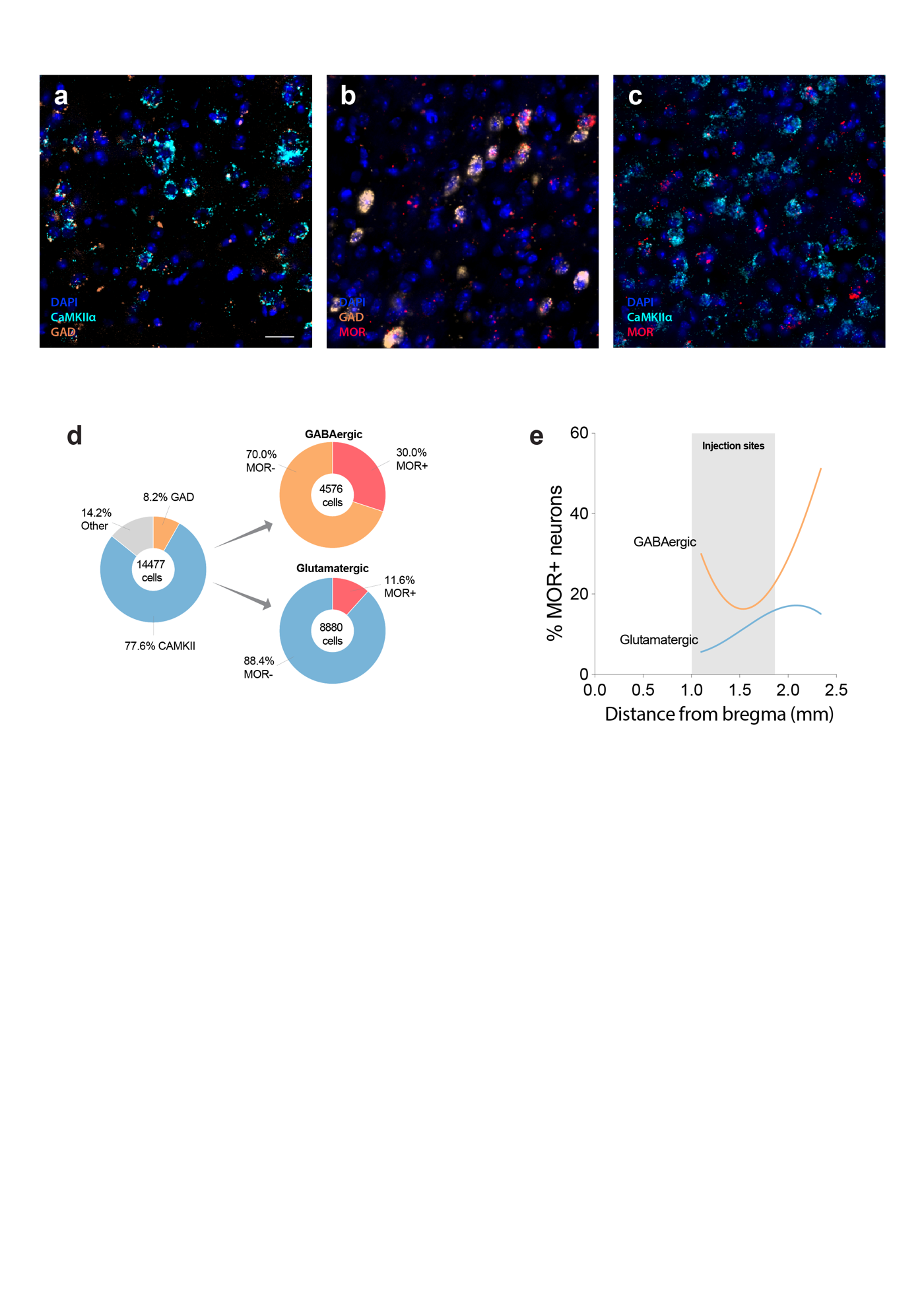

**Supplementary Figure 14. RNAscope validation of μ-opioid receptor expression in glutamatergic and GABAergic neurons in the anterior cingulate cortex.** **a–c** Representative RNAscope images illustrating detection of Oprm1 (MOR) transcripts in CaMKIIα-expressing glutamatergic neurons and GAD-expressing GABAergic neurons within the anterior cingulate cortex (ACC). Sections were labeled for CaMKIIα, GAD, and Oprm1 transcripts with DAPI marking nuclei. Example images show CaMKIIα and GAD expression (**a**), GAD and Oprm1 co-localization (**b**), and CaMKIIα and Oprm1 co-localization (**c**). Scale bar = 20 μm. **d** Quantification of neuronal populations identified by RNAscope in the ACC (~+1.5 mm from bregma). Among all detected neurons (n = 14,477 cells), the majority were CaMKIIα+ glutamatergic neurons (77.6%), with smaller proportions of GAD+ GABAergic neurons (8.2%) and other cell types (14.2%). Oprm1 transcripts were detected in 30.0% of GABAergic neurons (n = 4,576 cells) and 11.6% of glutamatergic neurons (n = 8,880 cells). **e** Spatial distribution of MOR^+^ neurons along the anterior–posterior axis of the ACC based on RNAscope quantification. The percentage of MOR^+^ neurons was higher among GABAergic neurons than glutamatergic neurons across the ACC. The shaded region indicates the ACC injection coordinates used for dermorphin-saporin and chemogenetic experiments in the main study. Cells were quantified using QuPath subcellular detection based on RNAscope puncta localized around DAPI-defined nuclei.

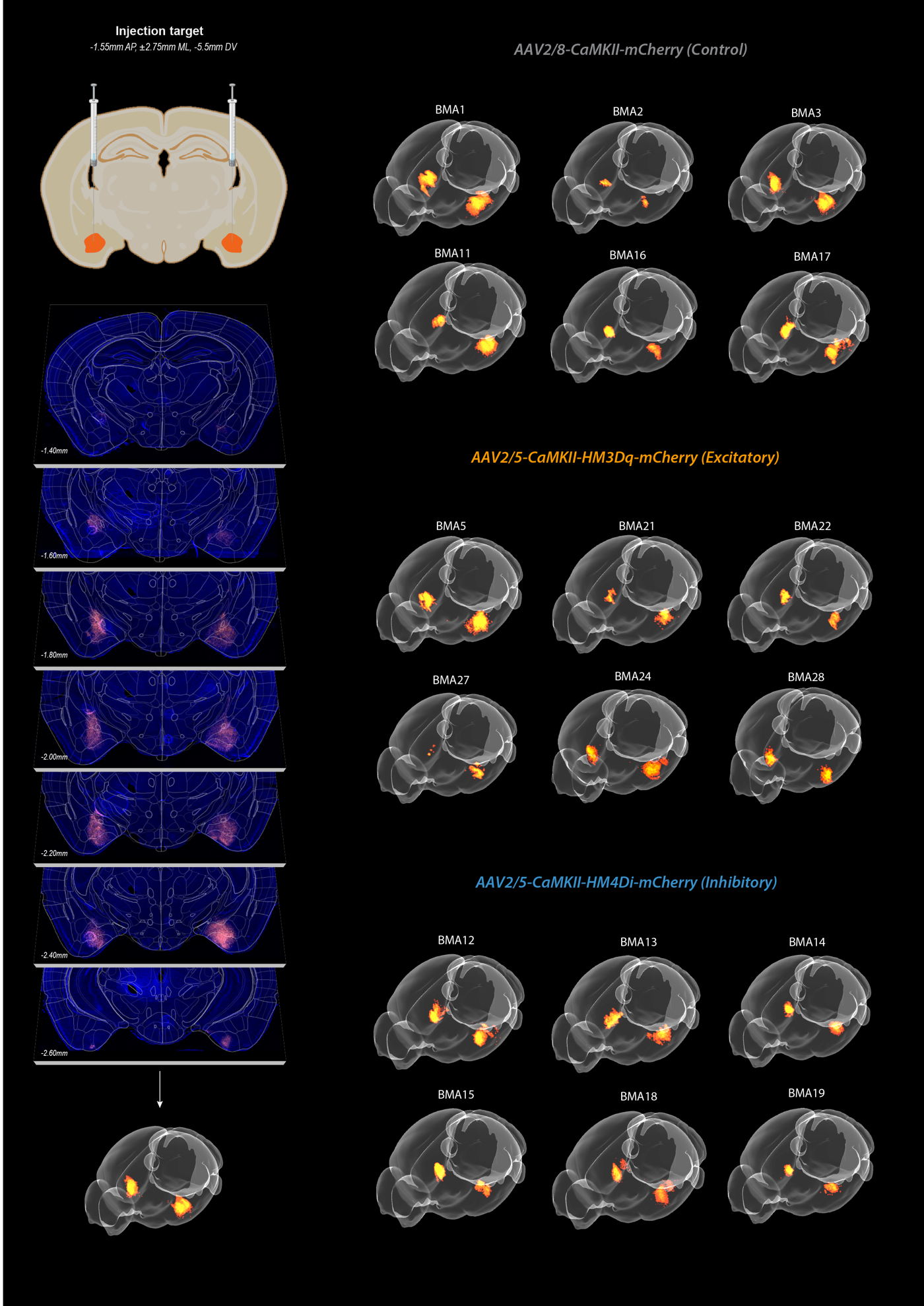

**Supplementary Figure 15. Verification of viral expression and injection targeting in the basomaedial amygdala (BMA).** (Top left) Schematic showing the intended injection coordinates targeting the BMA at -1.55mm AP, ±2.75mm ML, and –5.50mm DV relative to bregma. (Middle left) Histological verification of mCherry expression across coronal sections from a representative animal, shown at multiple anteroposterior levels spanning the BMA. (Bottom left) 3D reconstruction of mCherry expression from that animal using HERBS. (Right panels) Individual 3D reconstructions of mCherry expression for all included animals in each group are shown: AAV2/8-CaMKIIα-mCherry (Control, top), AAV2/5-CaMKIIα-hM3Dq-mCherry (Excitatory DREADD, middle), and AAV2/5-CaMKIIα-hM4Di-mCherry (Inhibitory DREADD, bottom). Each brain represents a distinct animal. Reconstructed volumes were aligned using HERBS software.

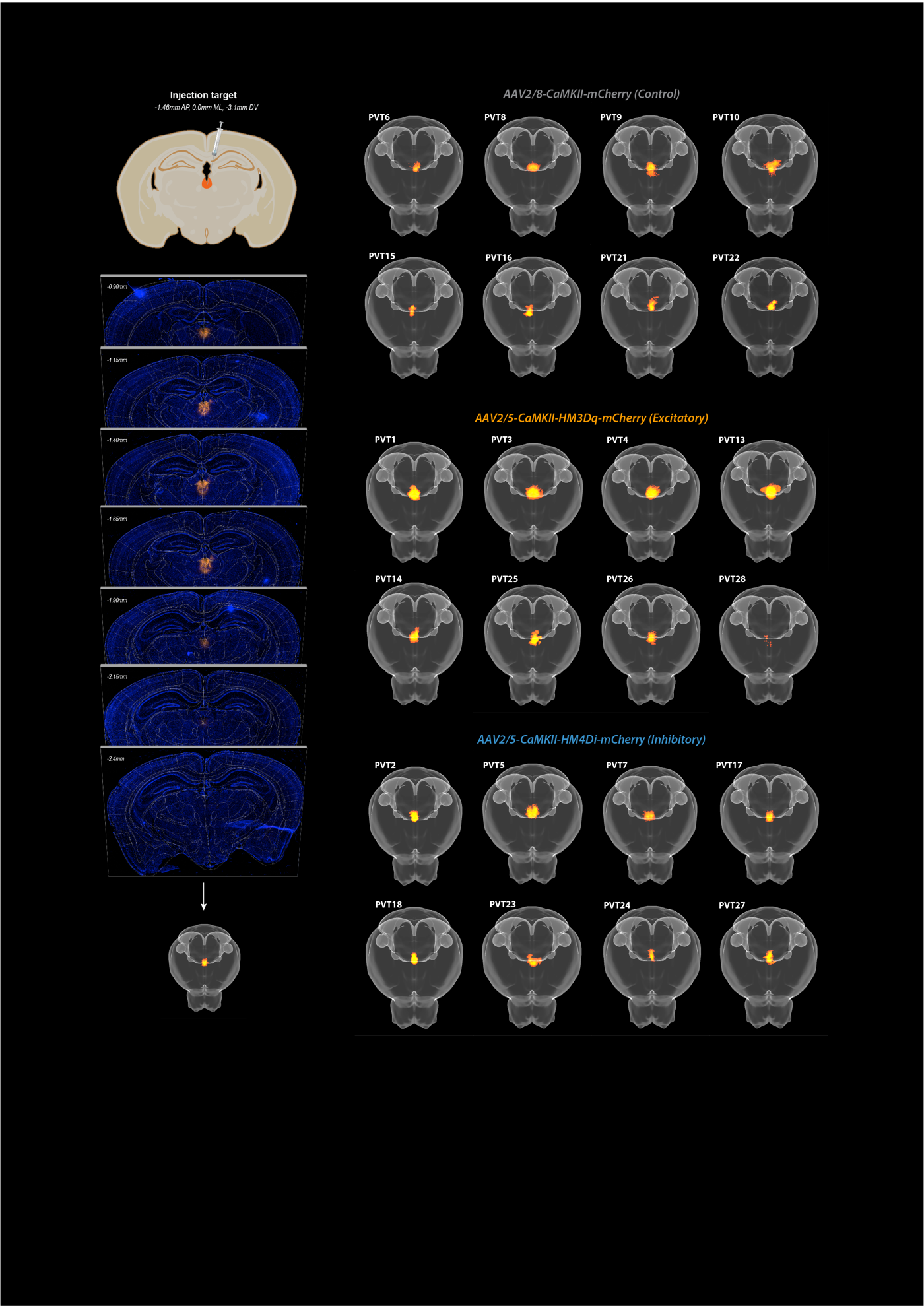

**Supplementary Figure 16. Verification of viral expression and injection targeting in the paraventricular thalamus (PVT).** (Top left) Schematic showing the intended injection coordinates targeting the PVT at –1.46mm AP, 0.0mm ML, and –3.1mm DV relative to bregma. (Middle left) Histological verification of mCherry expression across coronal sections from a representative animal, shown at multiple anteroposterior levels spanning the PVT. (Bottom left) 3D reconstruction of mCherry expression from that animal using HERBS. (Right panels) Individual 3D reconstructions of mCherry expression for all included animals in each group are shown: AAV2/8-CaMKIIα-mCherry (Control, top), AAV2/5-CaMKIIα-hM3Dq-mCherry (Excitatory DREADD, middle), and AAV2/5-CaMKIIα-hM4Di-mCherry (Inhibitory DREADD, bottom). Each brain represents a distinct animal. Reconstructed volumes were aligned using HERBS software.
