## Supplementary Table 1 for "Whole-brain analyses identify anterior cingulate μ-opioid signaling as a critical mediator of placebo analgesia in neuropathic pain"

### Supplementary Table1: Regions of Interest (ROIs) abbreviations list.

| **ROI Abbreviation** | **ROI Full Name** | **Regional Classification** |
| --- | --- | --- |
| mOFC | Medial Orbitofrontal Cortex | Cortical regions |
| vlOFC | Ventrolateral Orbitofrontal Cortex | Cortical regions |
| lOFC | Lateral Orbitofrontal Cortex | Cortical regions |
| IL(2/3) | Infralimbic Cortex Layer 2/3 | Cortical regions |
| IL(5/6) | Infralimbic Cortex Layer 5/6 | Cortical regions |
| PL(2/3) | Prelimbic Cortex Layer 2/3 | Cortical regions |
| PL(5/6) | Prelimbic Cortex Layer 5/6 | Cortical regions |
| dACC(2/3) | Dorsal Anterior Cingulate Cortex Layer 2/3 | Cortical regions |
| dACC(5/6) | Dorsal Anterior Cingulate Cortex Layer 5/6 | Cortical regions |
| vACC(2/3) | Ventral Anterior Cingulate Cortex Layer 2/3 | Cortical regions |
| vACC(5/6) | Ventral Anterior Cingulate Cortex Layer 5/6 | Cortical regions |
| SS Lower | Primary Somatosensory Cortex Lower Layers | Cortical regions |
| SS Upper | Primary Somatosensory Cortex Upper Layers | Cortical regions |
| dINS(2/3) | Dorsal Insular Cortex Layer 2/3 | Cortical regions |
| dINS(5/6) | Dorsal Insular Cortex Layer 5/6 | Cortical regions |
| vINS(2/3) | Ventral Insular Cortex Layer 2/3 | Cortical regions |
| vINS(5/6) | Ventral Insular Cortex Layer 5/6 | Cortical regions |
| NAc-C | Nucleus Accumbens Core | Accumbens |
| NAc-S | Nucleus Accumbens Shell | Accumbens |
| CA1 | Cornu Ammonis 1 | Hippocampus |
| CA2 | Cornu Ammonis 2 | Hippocampus |
| CA3 | Cornu Ammonis 3 | Hippocampus |
| DG | Dentate Gyrus | Hippocampus |
| BMA | Basomedial Amygdala | Amygdala |
| BLA | Basolateral Amygdala | Amygdala |
| LA | Lateral Amygdala | Amygdala |
| CeA | Central Amygdala | Amygdala |
| MeAv | Medial Amygdala, Anteroventral Part | Amygdala |
| MeAd | Medial Amygdala, Anterodorsal Part | Amygdala |
| CoA | Cortical Amygdala | Amygdala |
| IA | Interanterodorsal Thalamic Nucleus | Thalamus |
| LHb | Lateral Habenula | Thalamus |
| PVT | Paraventricular Thalamic Nucleus | Thalamus |
| CM | Central Medial Thalamic Nucleus | Thalamus |
| CL | Central Lateral Thalamic Nucleus | Thalamus |
| IMD | Intermediodorsal Thalamic Nucleus | Thalamus |
| RH | Rhomboid Thalamic Nucleus | Thalamus |
| RE | Reuniens Thalamic Nucleus | Thalamus |
| MD | Mediodorsal Thalamic Nucleus | Thalamus |
| RT | Reticular Thalamic Nucleus | Thalamus |
| VPL | Ventral Posterolateral Thalamic Nucleus | Thalamus |
| PreOp | Preoptic Area | Hypothalamus |
| ZI | Zona Incerta | Hypothalamus |
| LHA | Lateral Hypothalamic Area | Hypothalamus |
| Me | Median Eminence | Hypothalamus |
| DMH | Dorsomedial Hypothalamus | Hypothalamus |
| VMH | Ventromedial Hypothalamus | Hypothalamus |
| Posterior | Posterior Hypothalamus | Hypothalamus |
| LC | Locus Coeruleus | Brainstem |
| VTA | Ventral Tegmental Area | Brainstem |
| dmPAG | Dorsomedial Periaqueductal Gray | Brainstem |
| dlPAG | Dorsolateral Periaqueductal Gray | Brainstem |
| lPAG | Lateral Periaqueductal Gray | Brainstem |
| vlPAG | Ventrolateral Periaqueductal Gray | Brainstem |
| DR | Dorsal Raphe Nucleus | Brainstem |
| RVM | Rostral Ventromedial Medulla | Brainstem |
