## Supplementary Table 2 for "Whole-brain analyses identify anterior cingulate μ-opioid signaling as a critical mediator of placebo analgesia in neuropathic pain"

### Supplementary Table2: Distribution of strongest functional connectivity edges around anatomical class

| **Connection class** | **Natural History (%)** | **Placebo (%)** |
| --- | --- | --- |
| Cortex-cortex | 8 | 39 |
| Cortex-limbic | 9 | 25 |
| Cortex–thalamus | 6 | 4 |
| Cortex–hypothalamus | 2 | 0 |
| Cortex–brainstem | 5 | 5 |
| Limbic–limbic | 5 | 9 |
| Limbic–thalamus | 11 | 4 |
| Limbic–hypothalamus | 4 | 2 |
| Limbic–brainstem | 8 | 6 |
| Thalamus–thalamus | 9 | 1 |
| Thalamus–hypothalamus | 7 | 0 |
| Thalamus–brainstem | 16 | 1 |
| Hypothalamus–hypothalamus | 1 | 2 |
| Hypothalamus–brainstem | 3 | 0 |
| Brainstem–brainstem | 6 | 2 |
| **Summary classification** |  |  |
| Cortical-cortical | 8 | 39 |
| Cortical-subcortical | 22 | 34 |
| Subcortical-subcortical | 70 | 27 |

Values indicate the percentage of edges within the top-100 strongest functional connections for each condition. Cortical regions included cortical ROIs, whereas subcortical regions included limbic, thalamic, hypothalamic, and brainstem structures.
